# ITC and SPR Analysis Using Dynamic Approach

**DOI:** 10.1101/788075

**Authors:** Ganesh Kumar Krishnamoorthy, Prashanth Alluvada, Shahul Hameed, Timothy Kwa, Janarthanan Krishnamoorthy

**Affiliations:** Curtiss-Wright Avionics and Electronics, Dublin 14, Ireland; Department of Bio-medical Engineering, Jimma Institute of Technology, Jimma University, Ethiopia; Wise corner Pvt Ltd, Chennai, Tamilnadu, India; #05-34, Blk 8, Holland Ave, Singapore

**Keywords:** Dynamic approach, ITC, SPR, Instrument response, Equivalent binding, Sequential binding, Aggregation model, RNASE, hBCL_XL_, BH3I-1

## Abstract

Biophysical techniques such as Isothermal Calorimetry (ITC) and Surface Plasmon Resonance (SPR) are routinely used to ascertain the global binding mechanisms of protein-protein or protein-ligand interaction. Recently, Dumas etal, have explicitly modelled the instrument response of the ligand dilution and analysed the ITC thermogram to obtain kinetic rate constants. Adopting a similar approach, we have integrated the dynamic instrument response with the binding mechanism to simulate the ITC profiles of equivalent and independent binding sites, equivalent and sequential binding sites and aggregating systems. The results were benchmarked against the standard commercial software Origin-ITC. Further, the experimental ITC chromatograms of 2’-CMP + RNASE and BH3I-1 + hBCL_XL_ interactions were analysed and shown to be comparable with that of the conventional analysis. Dynamic approach was applied to simulate the SPR profiles of a two-state model, and could reproduce the experimental profile accurately.

## 1. Introduction

Biophysical techniques such as Isothermal Calorimetry (ITC), and Surface Plasmon Resonance are used to thermodynamically and kinetically characterize the binding mechanism of the protein-ligand or protein-protein interactions, respectively [1,2,3,4]. ITC measures the heat released or absorbed during the protein-ligand interactions [1,2,3,5,6], whereas, SPR measures the change in reflective angle of the incident light caused by surface waves called ‘Plasmon Polaritons’. Since plasmon polaritons, are sensitive to binding events occurring on a surface, it proportionally affects the angle of the reflected light [7,8].

In ITC experiments, the thermogram with asymmetric Gaussian-like peaks are numerically integrated and normalized with respect to the titrated ligand to obtain ‘Normalized Delta Heat’ data (NDH) [9,10]. Assuming a kinetic model, NDH data is then analysed to obtain the stoichiometry, binding equilibrium, and enthalpy constants. On the other hand, the thermogram (time domain data without integrating the peaks) can be directly analysed as shown by Dumas etal, to obtain the kinetic rate constants [11]. Interestingly, the delay in ligand dilution after each injection and the heat released or absorbed due to binding events were modelled as a first order ‘Instrument response’. While doing so, the Instrument response is modelled independent of the kinetic binding mechanism. In this current work, we have modelled the instrument response and incorporated it within the kinetic framework.

In a conventional SPR experiment, the Response Unit (RU) of the SPR sensogram exhibits, three distinct phases such as association, dissociation and regeneration [7]. These phases are either analysed separately/peicewise or in an integrated manner to obtain the kinetic rate constants. Dynamic approach, as applied to ITC can also be easily extended to model the SPR profiles of different binding mechanisms [12,13,14,15]. Yet again the advantage of dynamic approach is that the instrument response can be seamlessly integrated within the kinetic framework of the binding mechanism thereby simplifying the data analysis. A representative of different kinds of binding mechanisms such as, single set of equivalent sites, two sets of equivalents sites with sequential binding mode, two sets of equivalents sites with parallel binding mode, and aggregation have been considered here [16,17,18,19,20,21,22,23,24,25,26,27]. The simulation of ITC profiles for all these mechanisms were realised and found to be consistent with that of the previous reports [9]. Further, we have also analysed the experimental ITC data of 2’-CMP + RNASE and BH3I-1 + hBCL_XL_, and determined the kinetic and thermodynamic parameters of the binding process. The results were consistent with the earlier reports [28,29]. Dynamic approach based simulation of the SPR profiles for a single binding site also yielded accurate profiles similar to reported experimental profiles.

## 2. Theory

Protien-ligand/ Protien-protein binding mechanism can be as simple as a single site binding or as complex as multi step sequential binding. Previous modelling approaches have taken into account the instrument repsonse, independent of the kinetic mechanism; wherein, the correction for ligand dilution and heat detection were made outside of the dynamics of the binding mechanism. Firstly, a detailed comparison of different modelling approaches, such as lumped modelling, sequential kinetic modelling, and the current parallel kinetic modelling, using a simple single site binding mechanism, is provided, Sec (2. 1) & SI(1.1 -1.4); Fig (1A-H). Secondly, how dynamic approach can be applied to model complex cases such as a single set of equivalent binding sites Sec (2.2.1), two sets of equivalent sites with sequential Sec (2.2.2) or parallel binding modes SI (2.1), and aggregation SI (2.3) are presented Fig (2). Extension of this approach to SPR for a single binding case, with and without ligand leakage is detailed in SI (3), Fig (3).

**Figure 1.**
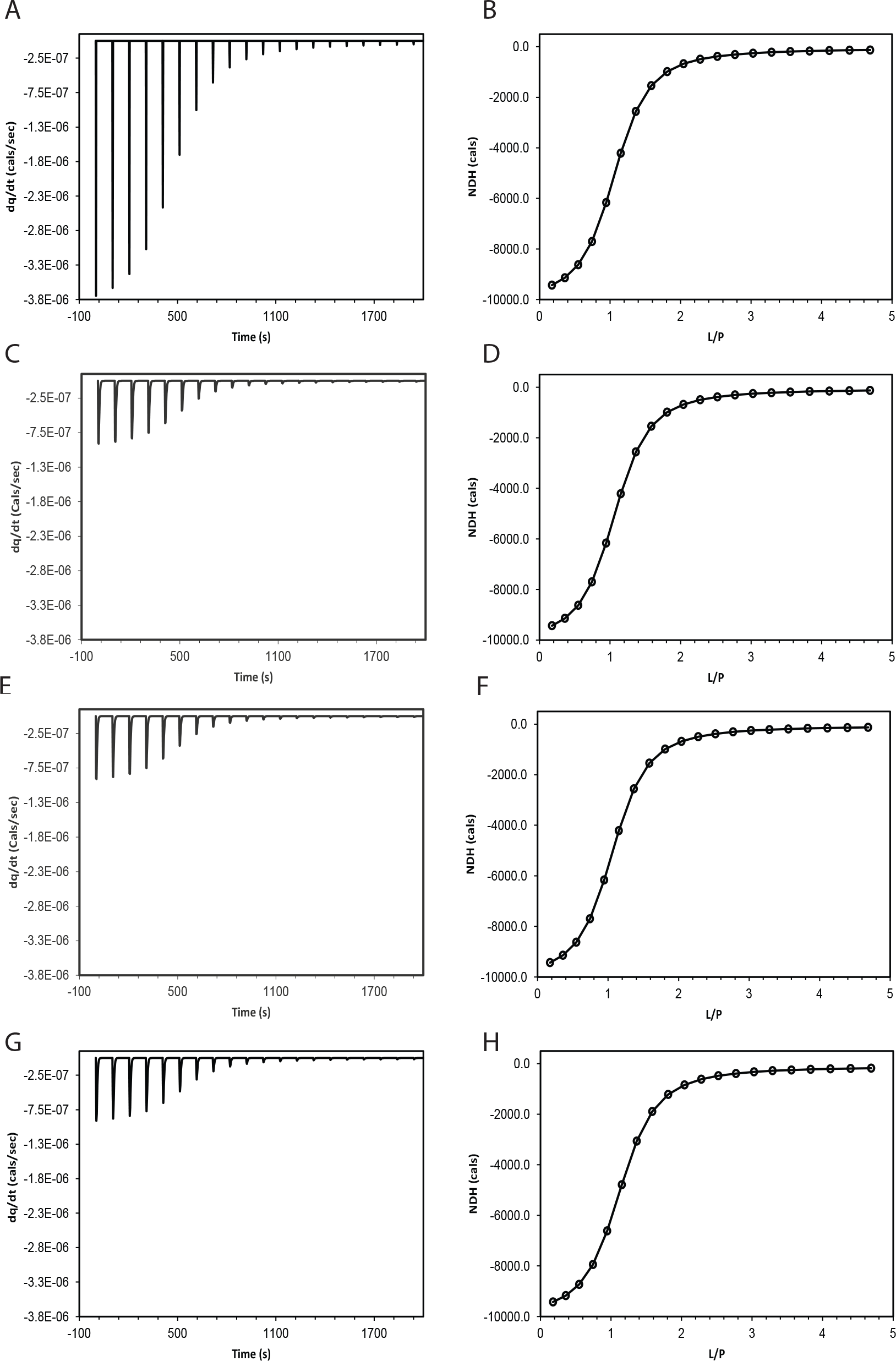
Comparison of the thermogram and the NDH data obtained for a single binding site mechanism through four different approaches. (A,B) without instrument response; (C,D) with instrument response calculated by lumped modelling; (E,F) with instrument response calculated by kinetic modelling in a sequential manner; (G,H) with instrument response calculated by kinetic modelling in a parallel manner.

**Figure 2.**
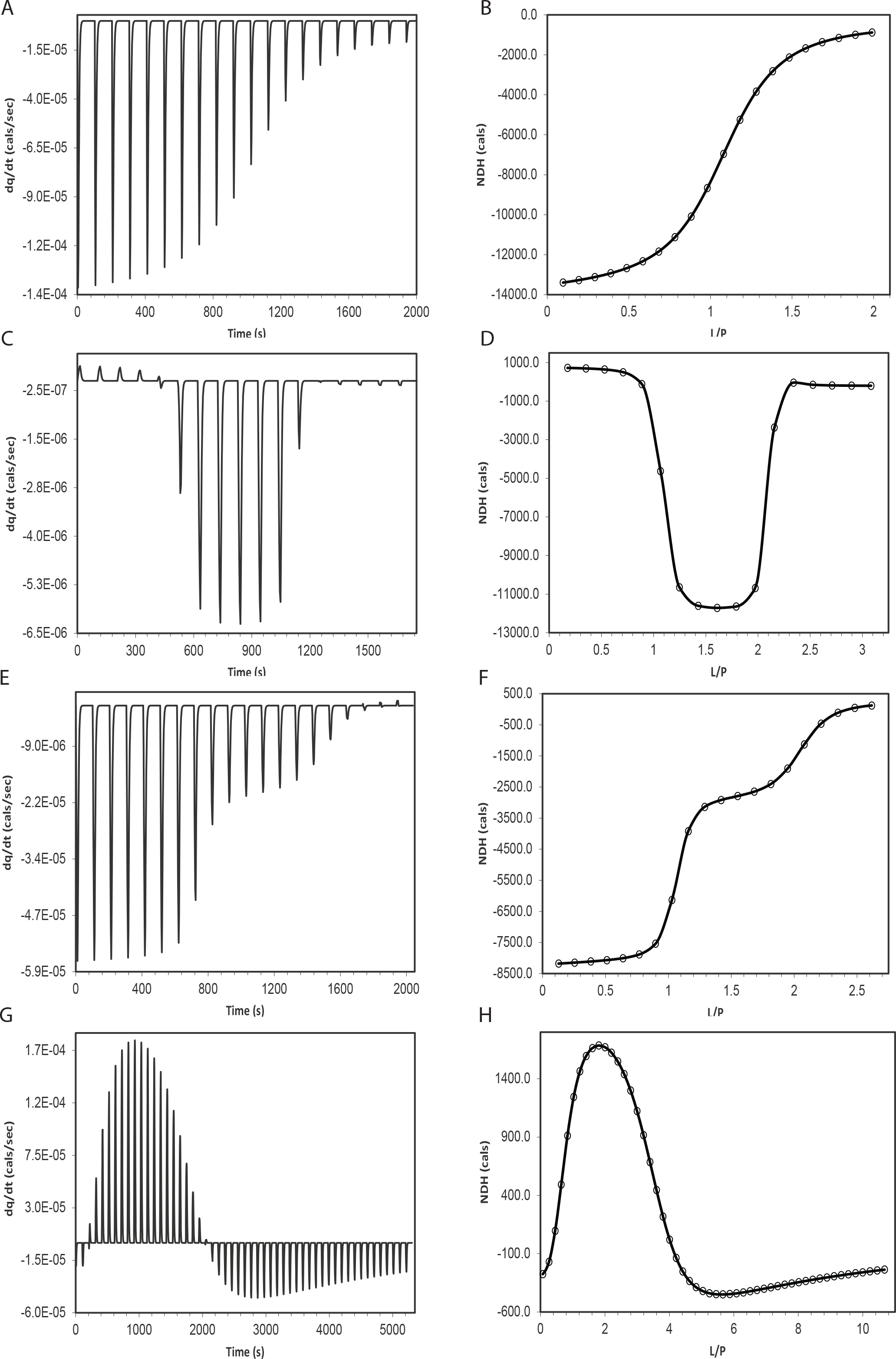
Simulation of the ITC thermogram and its corresponding NDH data for different binding mechanisms (A,B) M equivalent single site binding; (C,D) M, N, two equivalent independent/parallel binding sites (FEOTF54); (E,F) M, N, two equivalent sequential binding sites (PROTDB). (G,H) M, N, O, R, four equivalent sequential binding sites (PERSSON). In the NDH plots of B, D, F, H, the open circle represents the standard experimental data points as provided in Origin-ITC, and the smooth line represents the NDH data obtained by integrating the simulated thermogram shown in A, C, E, G, respectively.

**Figure 3.**
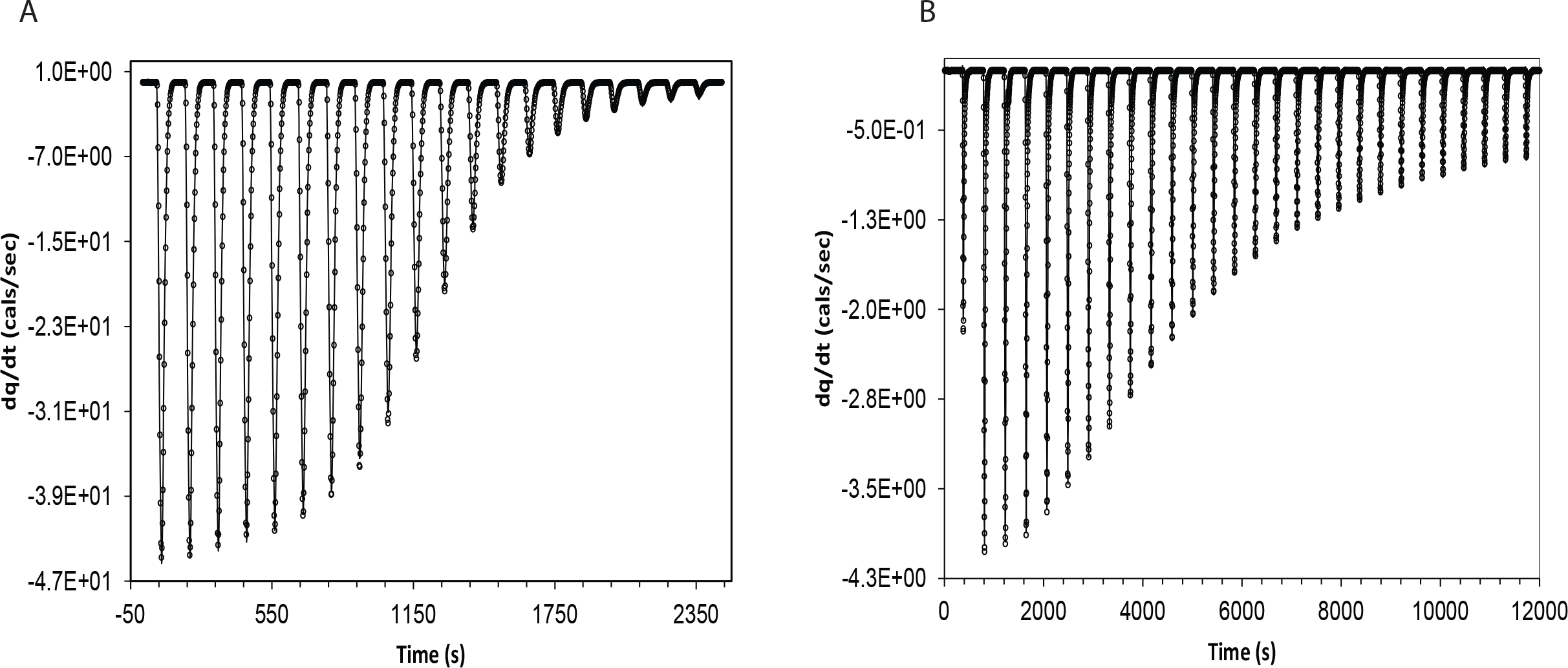
The experimental data and its model fit for (A) 2’-CMP + RNASE system using M equivalent single site binding, (B) BH3I-1 + hBCL_XL_ using M, N, two sequential binding sites.

### 2.1. Dynamic model with instrument response

In this section, we outline the kinetic based modelling approach, which incorporates instrument response within the binding mechanism. The kinetic mechanism can be proposed as follows,

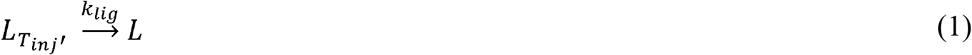

Where, 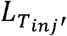, is the concentration of the injected ligand and *L*, is the concentration of the ligand available for binding after dilution. Eqn (1) accounts for the instrument response of the ligand dilution from the center of the ‘cell’ into the bulk of the solution.

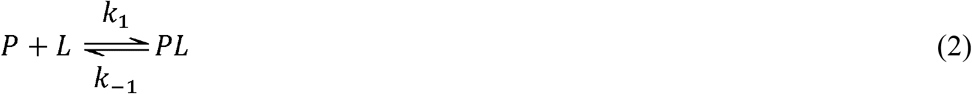

Eqn (2) accounts for the binding mechanism of the available ligand, *L*, with the protein, *P*, to form the complex, *PL* and *k*_1_, *k*_−1_, are the forward and reverse kinetic rate constants of the complex formation, respectively.

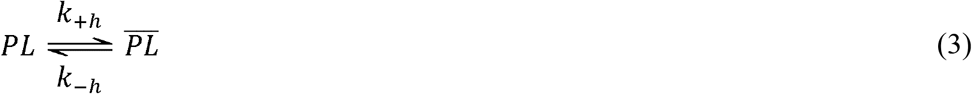

Eqn (3) accounts for the instrument response related to the heat released or absorbed caused by binding process. Here, the heat response rate (*k*_+*h*_) is related to the time delay 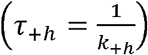, observed between the instance of heat generation and the instance of heat signal detected by the sensor/detector. As a simplification, we will always assume that, (*k*_+*h*_ = *k*_−*h*_), in the following discussion. The detected heat is represented in terms of ‘power’ in the thermogram.We have equated the ‘power’ pertaining to the binding process to be proportional to rate of change of 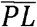, rather than *PL* itself. Following the same line of thought, Eqn (4) accounts for the instrument response associated with heat of ligand dilution. Here, the ‘power’ due to dilution is made proportional to the rate of change of 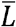, rather than *L* itself.

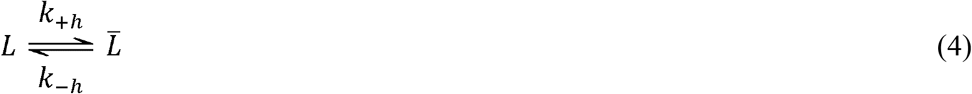

We have made an assumption here that the heat of dilution is primarily arising from the ligand rather than any buffer component or co-solvents. This assumption is valid because, in most of the experiments, the concentrations of the buffer and the co-solvents are maintained identical in both the cell and the injected ligand solutions, so as to avoid the heat of dilution due to buffers and co-solvents. The rate equations for the above kinetic mechanism can be written as Eqn (5–10),

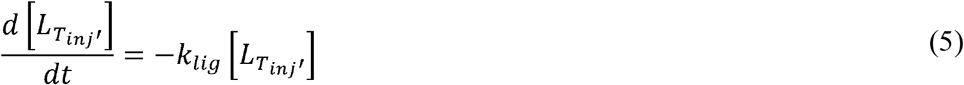

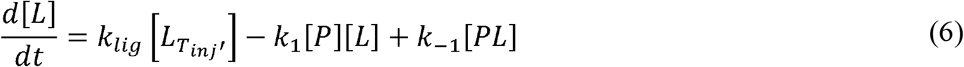

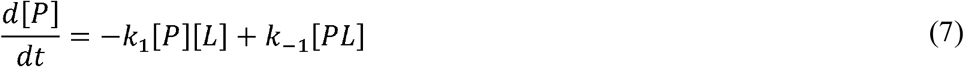

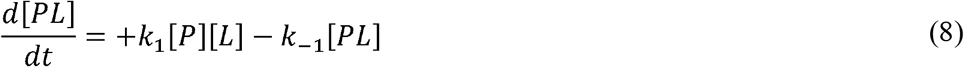

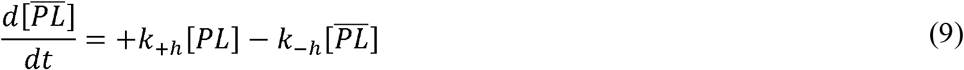

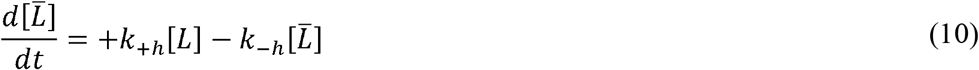

The above set of differential equations, Eqns (5–10), can be solved simultaneously to obtain, 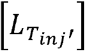, [*L*], [*P*], [*PL*],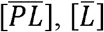 at different time points. Given the time profile of 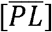, the time profile of the ‘power’ due to binding 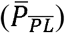 can be calculated as follows, Eqn (11),

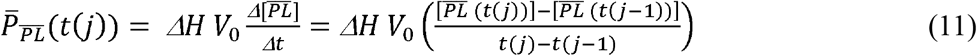

Where, 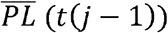 and 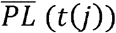 are the concentrations of complex, 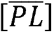 at *t*(*j* − 1) and *t*(*j*), respectively. Given the time profile of 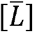, the time profile of the ‘power’ due to ligand dilution 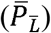 can be calculated as follows Eqn (12)

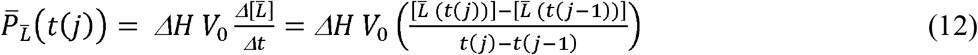

Where, 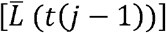 and 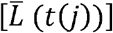 are concentrations of, 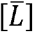 at *t*(*j* − 1) and *t*(*j*), respectively.

The simulated ITC chromatogram, is nothing but the sum of ‘power’ due to binding and dilution of ligand, 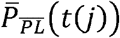 and 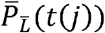. Comparison of four different approaches used to calculate the ITC time profile including the method described here SI (1.1-1.4) are shown in Fig (1 A-H). The parameters used to simulate these profiles are provided in Table SI 1.

### 2.2. Complex binding mechanisms

In this section, we provide the explicit kinetic mechanism for the complex binding cases such as,

1. *M* equivalent single site binding (E.g. RNAHH; Fig 2A, B).
2. *M*, *N* two sets of equivalent sites with sequential binding (E.g. PROTDB; Fig 2E, F). Other complex cases such as
3. *M*, *N*, two sets of equivalent sites and independent binding(E.g. FEOTF54; Fig 2C, D).
4. *M*, *N*, *O*, *R*, four sets of equivalent and sequential binding(E.g. PERSSON; Fig 2G, H).
5. Aggregation of *M* proteins.

are explained in the supplementary SI (2.1-2.3).

#### 2.2.1. *M* equivalent single binding site

In this mechanism, the ligand binds to a single site available in the protein, but the binding site can exist in ***M*** equivalent conformational forms. At the outset the ligand ‘available’ ***L*** for the binding process is modelled as given by Eqn(**1**). The kinetic mechanism of ***M*** equivalent sites binding can be outlined as in Eqn (**13**),

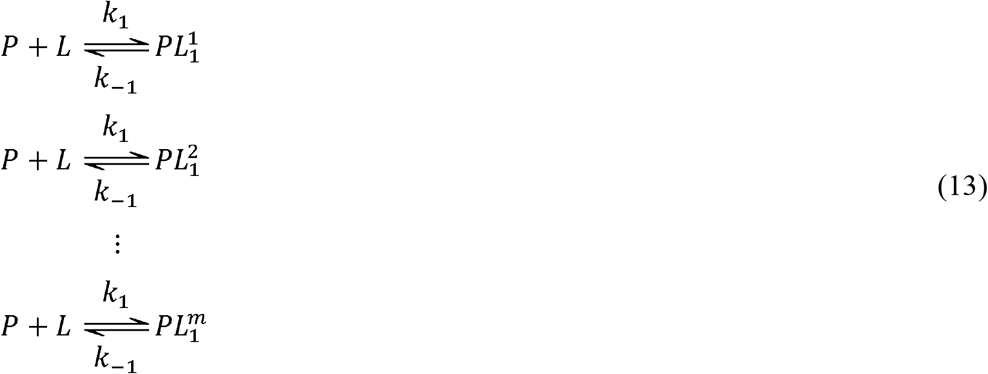

In the above representation of the complexes, 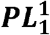 to 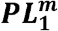, the superscript index denotes, **1** to ***m***, different equivalent sites; the subscript denotes the stoichiometry of ligand bound to the protein. The ‘power’ required to simulate the ITC profile, can be modelled as follows (Eqn (**14**)),

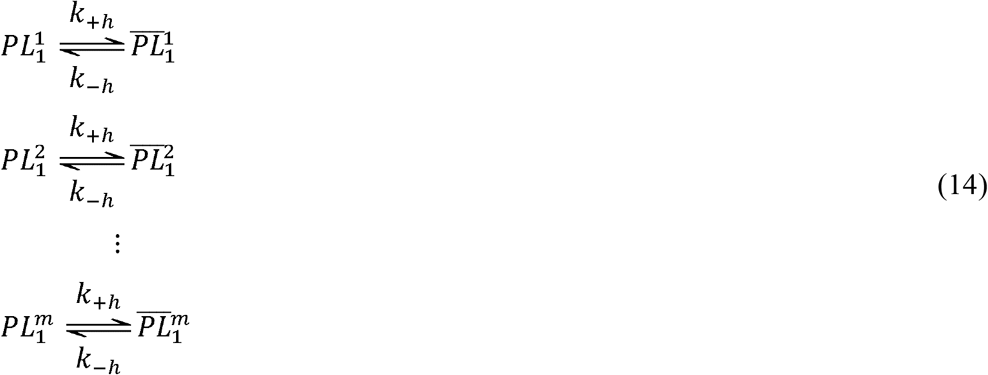

Where, ***k***_+*h*_, ***k***_−*h*_, are the forward and reverse heat response rate constants of the heat detector, respectively, and are assumed to be equal to each other. In the above Eqns(**14**), the 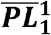 to 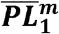, are the species, whose change in concentration generate the change in heat profile. While writing the mass balance for the total ligand or protein, we include only the species that are directly involved in the binding process (***P***, ***L***, 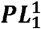 to 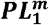), and exclude the species concerned with the instrument response such as 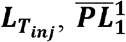 to 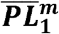. For example, the mass balance of the ligand would be, 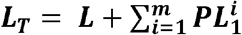 and for protein it would be, 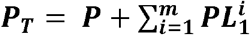. Additionally, we also consider a separate mass balance for the available and non-available ligand, that is independent of the binding process; 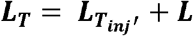. Based on the above proposed kinetic mechanism, the rate equations can be framed for each species and their time profile determined using numerical integration methods.

Before framing the rate equation, we make an assumption that the biding affinity of all the ‘***M***’ equivalent binding sites are equal. This also implies that the concentration of all the bound species such as 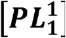 to 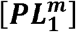 are all equal. If we represent, 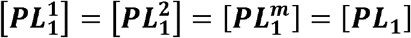, then the sum of all the bound complexes can be written as, 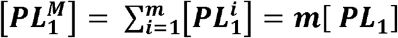. While writing the rate equations for the bound complex, only 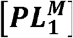 will be considered as a single entity, which represents the sum of all the equivalent bound forms.

The rate equation for the above kinetic mechanism, Eqns (1, 13, 14), can be written as Eqns (15–20),

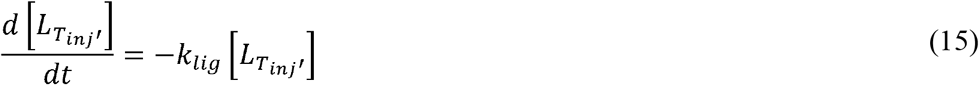

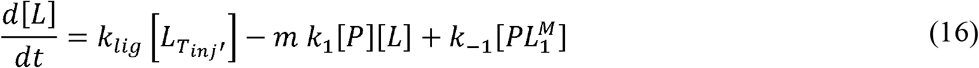

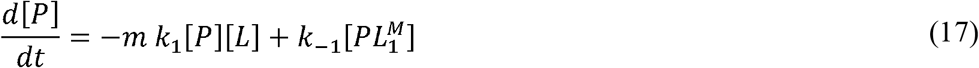

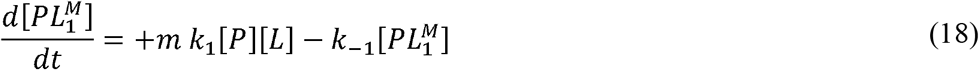

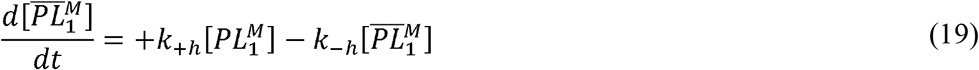

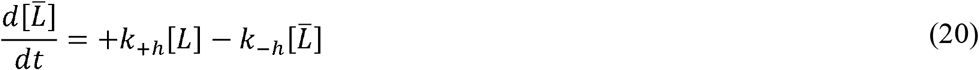

Where, ***m*** represents the total number of equivalent conformations, 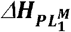, represents the enthalpy due to binding, Δ***H***_*dil*_, represents the enthalpy due to dilution, ***V***_0_, represents the total cell volume. The final ITC chromatogram 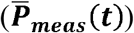 is the sum of ‘power’ due to binding (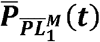: Eqn (21)) and ‘power’, due to ligand dilution (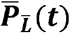: Eqn (22)) and is represented as, 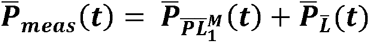, where,

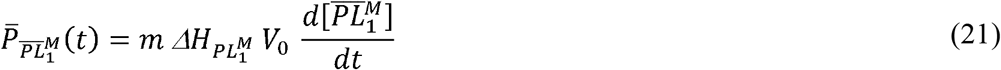

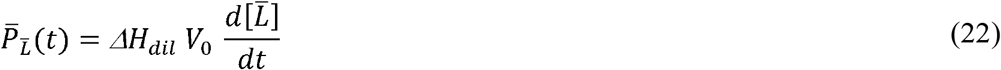

In the all the numerical simulations mentioned in this work we used the ‘difference’ form of the above equations Eqn(21, 22) rather than its differential form, as shown below Eqns (23, 24),

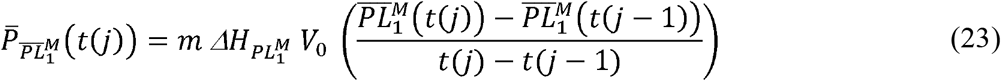

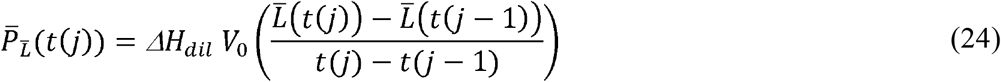

The simulation parameters and the corresponding ITC thermogram is presented in (Table 1) and Fig (2A,B).

**Table 1.** Parameters used to simulate the thermograms of different models using dynamic approach. These parameters are in turn obtained from fitting to equilibrium models available in Origin-ITC and are summarised in Table SI 5.

|  | RNAHH | FEOTF54 | PROTDB | PERSSON |
| --- | --- | --- | --- | --- |
| Model | One sites | Two sites<br>(independent) | Sequential binding<br>sites (2 sites) | Sequential binding sites<br>(4 sites) |
| Parameters |  |  |  |  |
| Kinetic<br>constants ($k$) | $5.59 \times 10^4 \text{ Hz}$<br>($k_1$)<br>1.0 ($\text{Hz}, k_{-1}$) | $1.18 \pm 0.40 \times 10^{10} \text{ Hz}$<br>($k_1$)<br>1.0 $\text{Hz}$ ($k_{-1}$)<br>$3.46 \pm 0.91 \times 10^6 \text{ Hz}$<br>($k_2$)<br>1.0 $\text{Hz}$ ($k_{-2}$) | $4.13 \times 10^7 \text{ Hz}$ ($k_1$)<br>1.0 $\text{Hz}$ ($k_{-1}$)<br>$1.40 \times 10^5 \text{ Hz}$ ($k_2$)<br>1.0 $\text{Hz}$ ($k_{-2}$) | $2.39 \times 10^3 \text{ Hz}$ ($k_1$)<br>1.0 $\text{Hz}$ ($k_{-1}$)<br>114 $\text{Hz}$ ($k_2$)<br>1.0 $\text{Hz}$ ($k_{-2}$)<br>$2.26 \times 10^3 \text{ Hz}$ ($k_3$)<br>1.0 $\text{Hz}$ ($k_{-3}$)<br>22.3 $\text{Hz}$ ($k_4$)<br>1.0 $\text{Hz}$ ($k_{-4}$) |
| Stoichiometric<br>constant | 1.02 ($m$ index) | 1.06 ($m$ index)<br>0.941 ($n$ index) | 1.0 ($m$ index; fixed)<br>1.0 ($n$ index; fixed) | 1.0 ($m$ index; fixed)<br>1.0 ($n$ index; fixed)<br>1.0 ($o$ index; fixed)<br>1.0 ($r$ index; fixed) |
| Thermodynamic<br>constants<br>($\Delta H$ cal/mol) | $-1.354 \times 10^4$<br>cal/mol ($\Delta H_{bind}$) | 767.3 cal/mol<br>($\Delta H_{bind_1}$)<br>$1.203 \times 10^4$ cal/mol<br>($\Delta H_{bind_2}$) | -8194 cal/mol<br>($\Delta H_{bind_1}$)<br>-3121 cal/mol<br>($\Delta H_{bind_2}$) | 314.8 cal/mol ($\Delta H_{bind_1}$)<br>4769 cal/mol ($\Delta H_{bind_2}$)<br>834.9 cal/mol ($\Delta H_{bind_3}$) |
| | | | | 6962 cal/mol ($\Delta H_{bind_4}$) |
| Injection, Cell parameters | 20 (Inj no)<br>4 μL(Inj vol)<br>1345 μL (Cell vol) | 17 (Inj no)<br>5 μL (Inj vol)<br>1411 μL (Cell vol) | 20 (Inj no)<br>4 μL(Inj vol)<br>1320 μL (Cell vol) | 52 (Inj no)<br>2 μL (1<sup>st</sup>), 5 μL, rest Inj vol)<br>1320 μL (Cell vol) |
| Instrument response | 0.33 Hz ($k_{lig}$)<br>0.33 Hz ($k_{+h}, k_{-h}$) | 0.33 Hz ($k_{lig}$)<br>0.33 Hz ($k_{+h}, k_{-h}$) | 0.33 Hz ($k_{lig}$)<br>0.33 Hz ($k_{+h}, k_{-h}$) | 0.33 Hz ($k_{lig}$)<br>0.33 Hz ($k_{+h}, k_{-h}$) |
| Other parameters | 651 μM (Prot)<br>21160 μM (Lig) | 31.4 μM (Prot)<br>1560 μM (Lig) | 494 μM (Prot)<br>20700 μM (Lig) | 6360 μM (Prot)<br>315000 μM (Lig) |
| Integration parameters per peak in the thermogram | 0 - 100 s (each inj)<br>50 (data points) | 0 - 100 s (each inj)<br>50 (data points) | 0 - 100 s (each inj)<br>50 (data points) | 0 - 100 s (each inj)<br>50 (data points) |

#### 2.2.2. *M*, *N*, two sets of equivalent binding sites with sequential binding

In this mechanism, there are two binding sites available for the ligand to bind to the protein in a sequential manner. The first binding site can exist in ***M*** equivalent conformational forms. The second binding site can exist ***N*** in equivalent conformational forms. At the outset the ‘ligand available’, ***L***, for the binding process is modelled using Eqn (1). The kinetic mechanism of the first binding step is represented as below, Eqn (25),

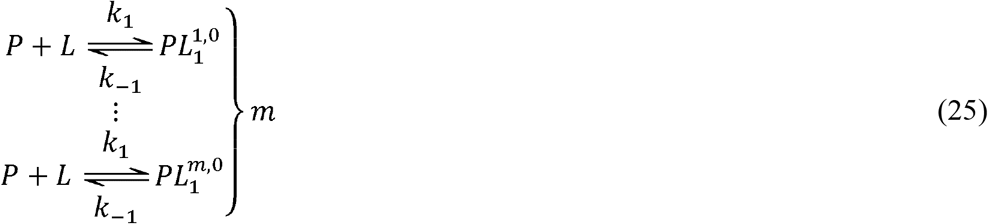

Where, ***k***_**1**_, ***k***_−**1**_, are the forward and reverse kinetic rate constants of the first binding process, respectively. In the above representation of the complexes, 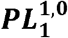 to 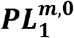, the superscript index has two place holders separated by comma. The first place holder is alloted for the first binding site and the corresponding index represent one of the ***m*** possible conformations or equivalent sites. The second place holder is alloted for the second binding site and the corresponding index represent one of the ***n*** possible conformations or equivalent sites. An index value of **0**, in any of the place holder indicates that the partcular binding process has not occurred yet. The subscript ‘1’, denotes that the stoichiometry of the ligand bound to the protein in all these complexes is ‘1’.

The second binding step which follows the first binding step, can be represented as follows, Eqn (26),

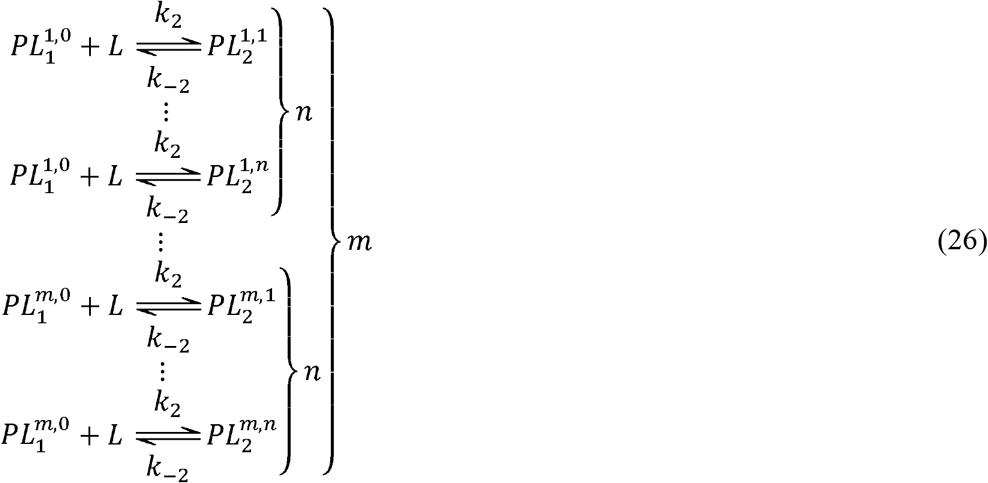

In the above representation of the complexes, 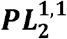 to 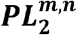, the superscript index denotes 1 to n equivalent sites, for the second binding process; the subscript ‘2’ denotes that the stoichiometry of the ligand bound to the protein in all these complexes is ‘2’.

As explained in the case of ***M*** equivalent single site binding, the concentrations of all the bound complexes after first binding steps can be represented as 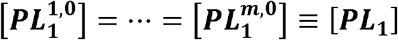, and the sum of all the bound complexes for the first binding process can be written as, 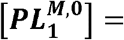 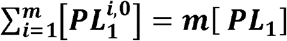. Similarly, the concentrations of all the bound complexes after second binding step can be represented as 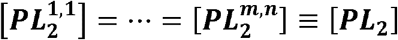, and the sum of all the bound complexes can be written as 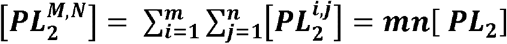. While writing the rate equations, for the bound complexes, only 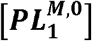 and 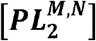 will be considered, which represent the sum of all the equivalent bound complex forms corresponding to the first and second binding processes, respectively. The heat response equilibrium can be written as Eqn (27),

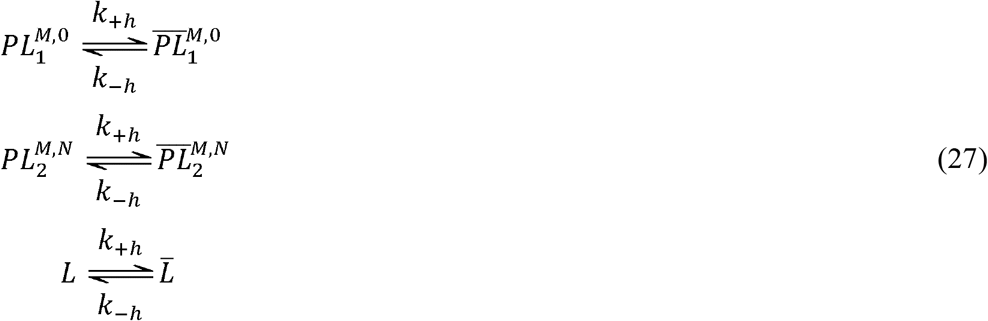

The rate equation for the above proposed kinetic mechanism can be written as, Eqns (28 - 35),

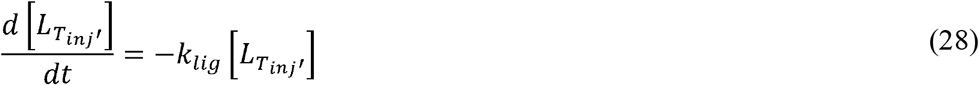

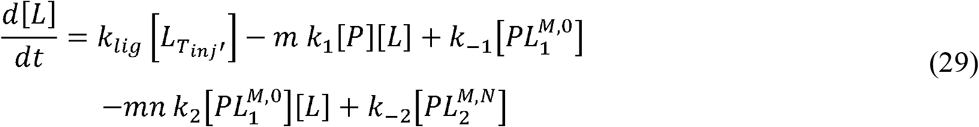

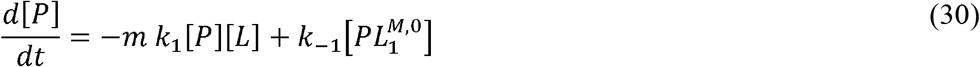

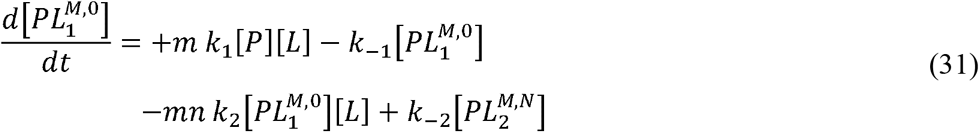

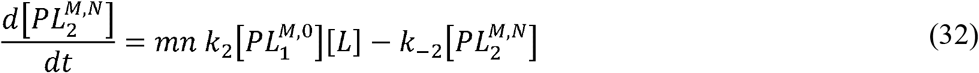

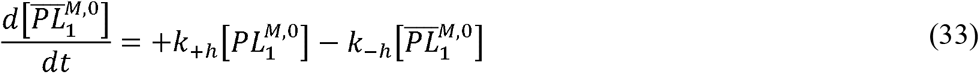

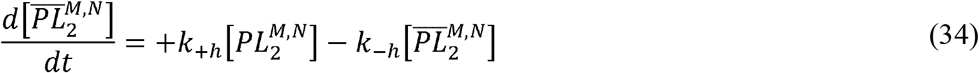

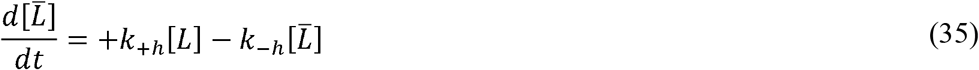

The final ITC chromatogram 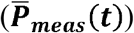 is the sum of the ‘power’ due to binding(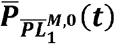: Eqn(36) and 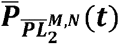: Eqn (37)) and ‘power’ due to ligand dilution (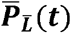: Eqn (38)) and is represented as 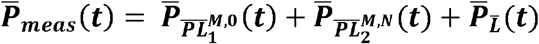, where,

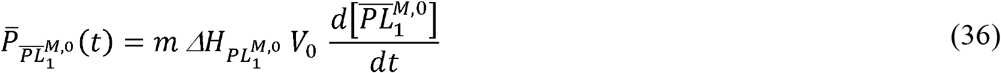

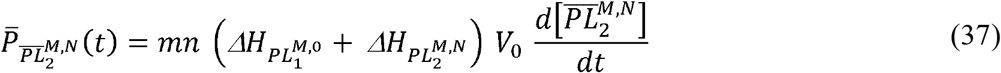

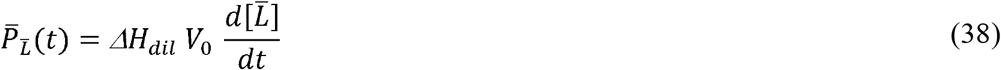

The simulation parameters and the corresponding ITC thermogram based on Eqns (28 – 38) is presented in (Table 1) and Fig (2E,F).

## 3. Results

Standard experimental data belonging to four different mechanisms such as (1) Single site (RNAHH) (2) Two sequential sites (PROTDB) (3) four sequential sites (PERSSON) and (4) two independent sites (FEOTF54) as provided in Origin-ITC software package were subjected to the conventional NDH based analysis [9,10]. The equilibrium constants and thermodynamic parameters obtained through optimization in Origin-ITC are tabulated in Table (SI 5). The same parameters were used to simulate the thermograms of RNAHH (Fig (2 A,B)), FEOTF54 (Fig (2 C,D)), PROTDB (Fig (2 E,F)) or PERSSON (Fig (2 G,H)) using dynamic models such as (1) M equivalent single site, (2) M, N equivalent parallel/independent binding sites and (3) M, N (or M, N, O, R) equivalent sequential sites (2 or 4 sites) respectively, Fig (2), Table (1). The NDH data resulted from the integration of the simulated thermograms were identical to that of the NDH values obtained from the experimental data. Additionally, simulations for protein dimerization and octamer formation were also performed with the aggregation parameters as tabulated in Table (SI 2).

The experimental ITC data of 2’-CMP + RNASE and BH3I-1 + hBclXL were analysed using M equivalent single site model and M, N Equivalent two site sequential model, respectively Table(2) & Fig (3A,B). The model fitted well to the experimental data and the results were comparable to that of the previously reported values [28,29]. In our current analysis, because of large residual values in initial fits for BH3I-1 + hBCL_XL_, we let the injection volumes to be varied during the optimization. While analysing, 2’-CMP + RNASE and BH3I-1 + hBCL_XL_, we fixed the protein, ligand and cell volumes as constants. Probability density based (PDF) based global sensitivity analysis on all fit parameters suggested that the model is equally sensitive to all the parameters used in the model, SI(4) & Fig (SI 2) [30,31].

**Table 2.** Fit parameters for 2’-CMP + RNASE and BH3I-1 + h BCL_XL_ based on dynamic approach.

|  | 2'-CMP + RNASE | BH3I-1 + hBclXL |
| --- | --- | --- |
| Model | M Equivalent single site model | M,N Equivalent two sequential model |
| Parameters |  |  |
| Kinetic rate constants ($k$) | $1.15 \pm 0.06 \times 10^3 \text{ Hz } (k_{on})$<br>$2.0 \pm 1.0 \times 10^2 \text{ Hz } (k_{off})$ | $3.09 \pm 0.72 \times 10^3 \text{ Hz } (k_{on_1})$<br>$45.0 \pm 4.0 \times 10^2 \text{ Hz } (k_{off_1})$<br>$2.56 \pm 1.12 \times 10^3 \text{ Hz } (k_{on_2})$<br>$1.39 \pm 0.25 \text{ Hz } (k_{off_2})$ |
| Equilibrium constants ($K_{eq}$) | $47.55 \pm 3.66 \times 10^3 \text{ } (K_1)$ | $6.82 \pm 1.67 \times 10^3 \text{ } (K_1)$<br>$1.84 \pm 0.81 \times 10^3 \text{ } (K_2)$ |
| Stoichiometric constant | $1.36 \pm 0.01 \text{ } (m \text{ index})$ | $1.75 \pm 0.26 \text{ } (m \text{ index})$<br>$2.55 \pm 0.54 \text{ } (n \text{ index})$ |
| Thermodynamic constants | $-1.02 \pm 0.00 \times 10^4 \text{ cal/mol } (\Delta H_{bind})$<br>$-1.34 \pm 0.37 \times 10^2 \text{ cal/mol } (\Delta H_{dil})$ | $-2.32 \pm 0.00 \times 10^4 \text{ cal/mol } (\Delta H_{bind_1})$<br>$-1.81 \pm 0.00 \times 10^4 \text{ cal/mol } (\Delta H_{bind_2})$<br>$4.99 \pm 1.14 \times 10^3 \text{ cal/mol } (\Delta H_{dil})$ |
| Instrument response | $7.11 \pm 1.29 \text{ Hz } (k_{lig})$<br>$6.47 \pm 1.29 \text{ Hz } (k_h)$ | $15.26 \pm 0.06 \times 10^3 \text{ Hz } (k_{lig})$<br>$3.86 \pm 0.12 \times 10^3 \text{ Hz } (k_h)$ |
| Corrections to the Injection vol | | $0.02 \pm 0.03 \text{ } \mu\text{L}$<br>$3.81 \pm 1.20 \text{ } \mu\text{L}$<br>$4.29 \pm 2.06 \text{ } \mu\text{L}$<br>$2.09 \pm 1.80 \text{ } \mu\text{L}$<br>$1.17 \pm 1.70 \text{ } \mu\text{L}$<br>$0.44 \pm 1.63 \text{ } \mu\text{L}$<br>$-0.08 \pm 1.58 \text{ } \mu\text{L}$<br>$-0.51 \pm 1.55 \text{ } \mu\text{L}$<br>$-0.92 \pm 1.51 \text{ } \mu\text{L}$<br>$-1.34 \pm 1.46 \text{ } \mu\text{L}$<br>$-1.76 \pm 1.41 \text{ } \mu\text{L}$<br>$-2.14 \pm 1.37 \text{ } \mu\text{L}$<br>$-2.45 \pm 1.33 \text{ } \mu\text{L}$<br>$-2.77 \pm 1.28 \text{ } \mu\text{L}$<br>$-3.06 \pm 1.24 \text{ } \mu\text{L}$<br>$-3.28 \pm 1.21 \text{ } \mu\text{L}$ |
|  |  | -3.51 ± 1.17 μL |
|  |  | -3.67 ± 1.15 μL |
|  |  | -3.72 ± 1.14 μL |
|  |  | -3.76 ± 1.13 μL |
|  |  | -3.82 ± 1.12 μL |
|  |  | -3.81 ± 1.12 μL |
|  |  | -3.80 ± 1.12 μL |
|  |  | -3.81 ± 1.11 μL |
|  |  | -3.58 ± 1.14 μL |
|  |  | -3.57 ± 1.13 μL |
|  |  | -3.35 ± 1.15 μL |
|  |  | -3.15 ± 1.17 μL |
|  |  | -2.79 ± 1.22 μL |

Simulation of SPR profile was carried out for a simple single site binding model (or two state model). The dynamic model was modified to account for the baseline anomalies often encountered in SPR profiles. By introducing a leakage factor in the ligand channel during dissociation, the concentration dependent residual changes in the baseline could be modelled accurately (Table SI 3) and Fig (4). The model used for ligand leakage (*L*(*t*)) during dissociation consisted of a constant part (basal value: *L*(0)) and an exponentially decaying function which is dependent on time (t); 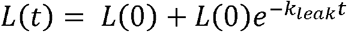; where, *k*_*leak*_, is the leakage rate. The simulation profiles were consistent with that of the earlier reported experimental profiles [7,8,32].

**Figure 4.**
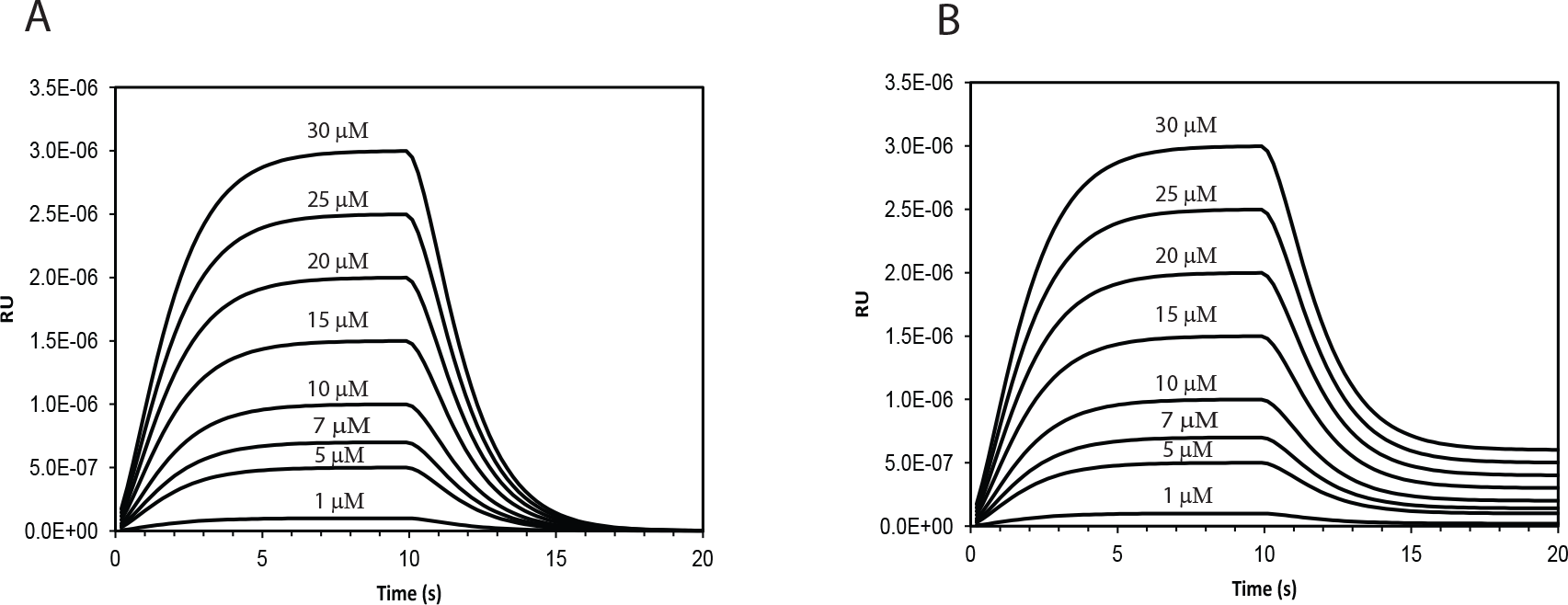
Simulation of the SPR sensogram using dynamic approach for a single binding site mechanism. (A) without any leakage of ligand during the dissociation phase (B) with leakage of ligand during the dissociation phase. The concentrations of the ligand used for each instance of the simulation is labelled above its respective traces.

## 4. Discussion

ITC thermogram is sensitive to various parameters such as protein concentration, ligand concentration, cell volume and injection volume [33]. The dilution effect of protein/ligand due to injected ligand, can be calculated as explained in SI (1.2.3). In real time situation, the dispersion of the highly concentrated ligand injected into the cell from the centre of the cell in to the bulk of the solution follows Fick’s law based diffusion formulated in the form of partial differential equation (PDE) Fig (5) [34,35]. In such a model, the concentration of ligand at each spatial location (X, Y, Z coordinates) varies with time, till the solution becomes homogeneous at equilibrium time point. This spatial and temporal dependence of the concentrations of the ligand and the protein can be simplified by taking into account the efficient stirring of the injector. If the mixing is effective, the spatial dependence can be ignored and the PDE reduces to an Ordinary differential equation (ODE) with dependence only on time. Under such circumstances, the dilution can be considered to be instantaneous, implying that equilibrium concentration is achieved instantaneously for the bulk solution, after the injection of ligand solution.

**Figure 5.**
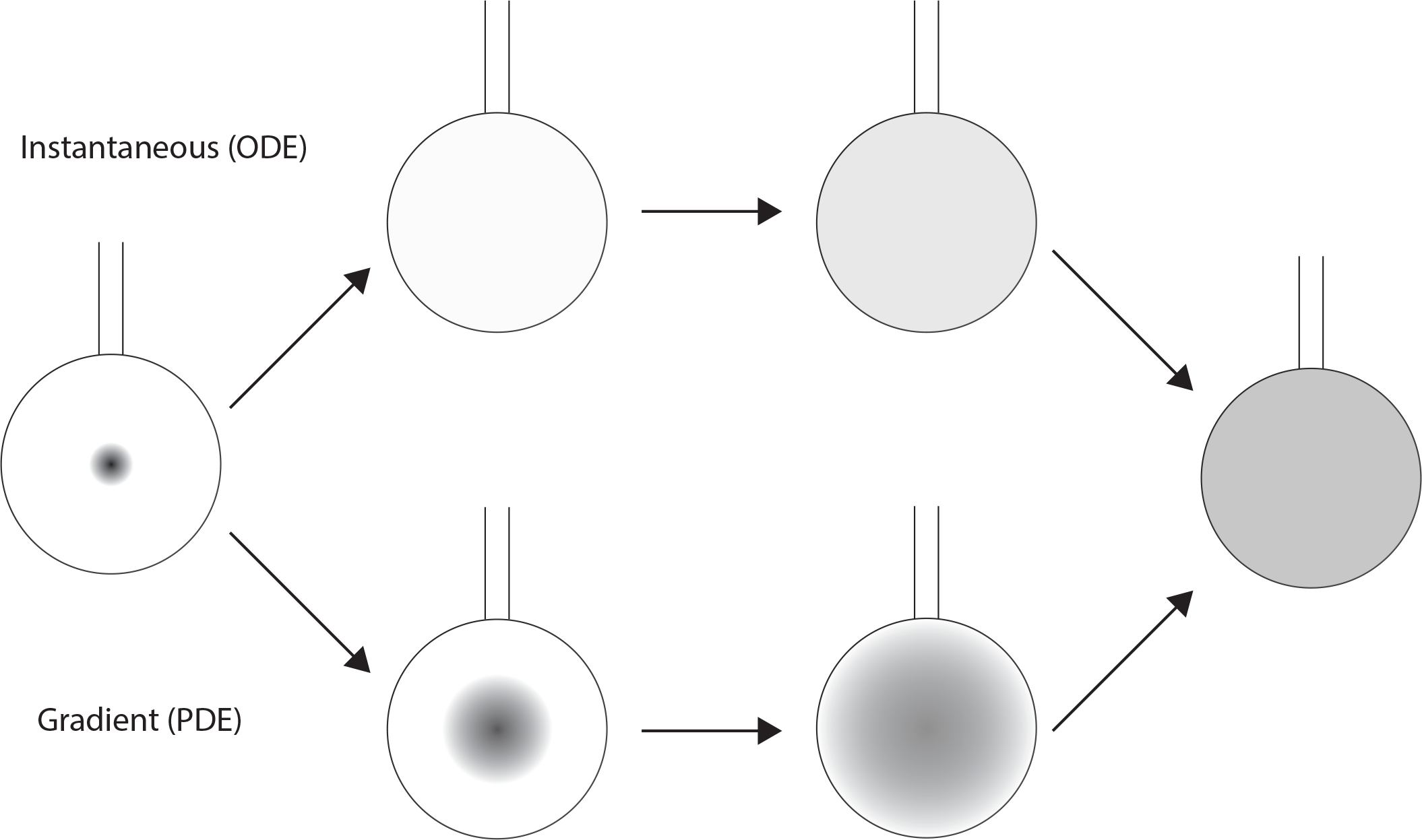
A comparison of the ligand dilution effect as addressed by ODE and PDE based dynamic modelling. The left and the right most figures represent the initial and final condition of the ligand concentrations immediately after injection and final equilibrium states, respectively. The darker shades represent higher concentration. In the upper scheme (ODE model), we assume an instantaneous mixing of sample being injected over discretized period of injection. Whereas, in the lower scheme (PDE model) we assume that the homogenization of the injected ligand is both time and spatial dependent.

Conventional analysis (or NDH analysis) of ITC data requires integration of each peaks in the thermogram and representing it as a function of protein to ligand ratio. In time domain analysis (Dynamic analysis), the thermogram is directly fitted to the model as explained in Sec 2.2.1 and 2.2.2, to obtain, thermodynamic and kinetic parameters. A closer inspection of the thermogram in Fig 1 A,C,E,G, clearly shows that though the time profile of with and without instrument response (left panel) differ significantly, the integrated data (right panel) is identical. This suggests that the heat energy represented by the area under the curve is conserved across time profiles of varied instrument response. In other words, NDH analysis is insensitive to instrument response seen with ligand dilution or heat detection.

In dynamic analysis, when an experimental thermogram is fitted against a kinetic model, the initial few data points may not fit well to the model. This is primarily due to the fact that, only a fraction of the data points contribute to the asymmetric Gaussian like thermogram profile, whereas, the rest of the data are baseline. Hence, during model fit, a weightage factor is calculated for each data point which is proportional to the difference of its value from the baseline. The model fit improved significantly, when the optimization was carried out with weightage factor. After optimization, probability density function (PDF) based sensitivity analysis was carried out on the fit parameters using SAFE Toolbox [30,31]. Basically, two PDF’s were calculated, namely, (1) conditional where one of the parameters is fixed and the rest of the parameters are varied and (2) unconditional, where all the parameters are varied. The difference between the conditional and unconditional PDF is a measure of sensitivity of that parameter to the given model. Based on this approach, all the optimized parameters were found to be invariably sensitive and significant to the models used in the analysis of RNASE and hBCLxl.

Dynamic analysis of 2’-CMP + RNASE experimental data yielded an equilibrium constant of 4.75×10^4^ and an enthalpy of −1×10^4^ Cals/mol. The fit parameters were comparable to that of the previous NDH based analysis (*K*_*eq*_:5.59 × 10^4^ M^−1^ and Δ*H*_*bind*_:−1.35 × 10^4^ Cals/mol) [9]. BH3I-1 + hBCL_XL_ on the other-hand was fitted to a *M*, *N* equivalent two site sequential binding site model. Previously, NDH based analysis carried out on the same data using a two-site sequential model with *M* = 1 and *N* = 1, yielded, *K*_1_ = 38 × 10^3^, *K*_2_ = 6 × 10^3^ M^−1^, which is comparable to the dynamic fit values of, *K*_1_ = 12 × 10^3^, *K*_2_ = 8 × 10^3^ M^−1^ [28,29]. Similarly the enthalpies from both the models were also comparable (NDH / (Dynamic analysis):- 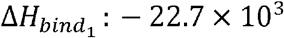, (−23.2 × 10^3^) Cals/mol and 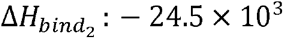, (−18.1 × 10^3^)) Cals/mol) [28,29].

Aggregation mechanisms are usually complex and difficult to model, due to the various binding modes available for the intra and inter molecular complex formation [25,27,36]. However, any complex aggregation mechanism can be modelled as a series of sequential binding mechanism, with several steps in between the monomer and the final aggregate polymer. In this study, we compared two aggregation cases, where a protein forms dimer (*M* = 2); and a octamer (*M* = 8). When the degree of polymerization for the protein increased from 2 to 8, distinct lag phase at the outset of the thermogram could be clearly seen [27]. In general, the ITC profile of the aggregation of a macromolecule in the presence of a ligand will exhibit an initial lag phase (flat line) correlated with the critical aggregation concentration (CAC), and is related to the non-cooperative binding. Subsequently, the co-operative phase steps in causing a rapid increase or decrease in Q value due to the intra-micellar aggregation. The higher order association between polymeric protein and polymeric ligand is characterized by the inter-micellar aggregation occurring at critical micellar concentration (CMC) [27]. Though our simulations are carried out only for the case of macromolecule aggregation, comparison could still be made with the first two phases of the experimental ITC profiles of SDS + PEG systems, exhibiting a characteristic lag and cooperative binding phases.

Conventional data analysis of SPR comprises of three distinct phases namely, association, dissociation and regeneration [7]. Since these three regimes are independent, these can be individually analysed using three separate analytical expressions. But dynamic approach integrate all these regimes seamlessly through a single set of ODE’s, which, not only accounts for kinetics but also the instrument response due to ligand dilution and detector. A common feature that often observed in the SPR profile is the concentration dependent residual baseline during dissociation/regeneration phase which had been modelled here successfully by considering ligand leakage.

## 5. Conclusion

Here, we have explicitly, modelled the general binding mechanisms often encountered in ITC experiments such as independent, sequential and aggregation. Using dynamic approach, the instrument response due to ligand dilution and heat detection were incorporated within the kinetic framework so as to simulate the experimental profiles accurately. Furthermore, the experimental thermogram of 2’-CMP + RNASE and BH3I-1 + hBCL_XL_ were analysed and the thermodynamic and kinetic parameters that are consistent with earlier studies were obtained. The same approach has been extended to simulate the SPR profiles of a single site binding mechanism.

## Supporting Information

Supporting Information contains explicit derivation of the models such as, thermogram without instrument response (Sec 1.1), with instrument response, (lumped model (Sec 1.2), kinetic sequential model (Sec 1.3) and kinetic parallel model (Sec 1.4)). Equivalence of all these models are also detailed (Sec 1.5). Mathematical expressions for the dilution effect of protein and ligand are provided (Sec 1.2.3). Complex binding models such as M,N, two equivalent and independent site model (Sec 2.1), M,N,O,R four state sequential model (Sec 2.2), M aggregation model (Sec 2.3) are provided. Dynamic model for simulating SPR profile is also provided (Sec 3.0). Details of the global sensitivity analysis for the fit parameters is provided (Sec 4.0). Matlab codes [37] to simulate ITC/SPR profiles and fit experimental ITC data will be provided in SI after the peer review.

## Acknowledgments

JK would like to thank Prof. Henry Mok and Prof. John Shriver for kindly providing the experimental data used in this work.

