## Supporting information for "ITC and SPR Analysis Using Dynamic Approach"

† Current Affiliation: Singapore Bio imaging Consortium, Astar Research Institutes, 11 Biopolis way, Singapore 138667.

### 1. Different modelling approaches to simulate ITC chromatogram profiles

Here, we use the term ‘sequential’ referring to the assumption that the ‘instrument response’ and the ‘binding kinetics’ occur sequentially and the calculations are carried out likewise. Whereas, the term ‘parallel’ refers to the simultaneous occurance of instrument repsonse and the binding events. In this section, we present four different approaches to model the ITC thermogram as listed below,

i. Model without any instrument response,

ii. Sequential model with instrument response uncoupled from binding mechanism (Lumped/Laplace approach),

iii. Sequential model with instrument response uncoupled from binding mechanism (kinetic approach),

iv. Parallel model with instrument response coupled with binding mechanism (kinetic approach)

#### 1. 1. Model without instrument response

In modelling without instrument response, we first present the conventional NDH based method, where the change in heat (DH or Delta H) is calculated by integrating the assymetric gaussian peak for each injection and normalizing (N) it with respect to the amount of ligand injected. Later on, the time domain analysis is detailed where, the thermogram itself is directly calculated.

#### 1.1.1. NDH based analysis

Consider a single site binding case where a protein, $\boldsymbol{P}$, binds to a ligand , $\boldsymbol{L}$ , to form the protein-ligand complex, $\boldsymbol{PL}$, Eqn (**1**).

$$P + L \underset{k_{-1}}{\overset{k_1}{\rightleftharpoons}} PL \tag{1}$$

The $\boldsymbol{k_1}$, $\boldsymbol{k_{-1}}$, are the forward and the reverse kinetic rate constants for the binding equilibrium, respectively. The equilibrium constants $\boldsymbol{K_1}$, can be defined as $\boldsymbol{K_1} = \frac{\boldsymbol{k_1}}{\boldsymbol{k_{-1}}}$ . Applying, Le-chaterlier's principle, we can write an algebraic expression for Equilibrium reaction Eqn (**2**), as,

$$K_1 = \frac{[PL]}{[P][L]} \tag{2}$$

The mass balance for total protein $\boldsymbol{P_T}$ , and total ligand $\boldsymbol{L_T}$ can written as Eqns (**3**, **4**),

$$P_T = [P] + [PL] \tag{3}$$

$$L_{\boldsymbol{T}} = [L] + [PL] \tag{4}$$

Using Eqn (**2**), Eqn (**3**), we can express $\boldsymbol{P}$, $\boldsymbol{PL}$, in terms of $\boldsymbol{L}$ as Eqn (**5**,**6**),

$$[P] = \frac{P_T}{1 + K_1[L]} \tag{5}$$

$$[PL] = \frac{P_T\, K_1[L]}{1 + K_1[L]} \tag{6}$$

Using Eqns (4, 6), we express, $\boldsymbol{L}$, as a polynomial with constants such as, $\boldsymbol{L_T}$, $\boldsymbol{P_T}$ and $\boldsymbol{K_1}$.

$$K_1\,[L]^2 + (K_1P_T - K_1L_T + 1)[L] - L_T = 0 \tag{7}$$

Among the two possible roots, the one that is positive, real (non-complex) and whose value is less than that of $\boldsymbol{L_T}$ is chosen as the feasible solution. Substituting the value of $\boldsymbol{L}$ back in to Eqn (**6**), yeilds the value of $\boldsymbol{PL}$.

In ITC experiments, the quantity of heat energy absorbed or released ($\varDelta\boldsymbol{Q}$) is proportional to the amount (mass) of the complex $\varDelta\boldsymbol{PL}$ formed in the solution after $\boldsymbol{i^{th}}$ injection of the ligand, Eqn (**8**)

$$\Delta Q(i) = \varDelta H\ (V_0\ \Delta[PL(i)]) \tag{8}$$

Where, $\varDelta\boldsymbol{H}$, is the change in enthalpy due to binding, $\boldsymbol{V_0}$, is the cell volume. $\varDelta[\boldsymbol{PL}(\boldsymbol{i})]$ is the change in the concentration of $[\boldsymbol{PL}]$, before and after $\boldsymbol{i^{th}}$ injection of ligand. Experimentally, $\Delta\boldsymbol{Q}(\boldsymbol{i})$, is nothing but the integrated area under the curve of the assymyetric gaussian peaks occuring in the thermogram. More explicitly $\Delta\boldsymbol{Q}(\boldsymbol{i})$ is given as Eqn (**9**),

$$Q_{Inj}(i) - Q_{Inj}(i-1) = \varDelta H\ V_0([\ PL(i)] - [\ PL(i-1)]) \tag{9}$$

Where, $\boldsymbol{Q_{Inj}}(\boldsymbol{i}-\boldsymbol{1})$, $\boldsymbol{Q_{Inj}}(\boldsymbol{i})$ are the total/cumulative heat released or absorbed prior to and after $\boldsymbol{i^{th}}$ injection of the ligand. $[\ \boldsymbol{PL}(\boldsymbol{i}-\boldsymbol{1})]$, $[\ \boldsymbol{PL}(\boldsymbol{i})]$, are the concentrations of $\boldsymbol{PL}$ prior to and after $\boldsymbol{i^{th}}$ injection of the ligand.

Usually, $\Delta\boldsymbol{Q}(\boldsymbol{i})$ is normalized ($\Delta\boldsymbol{Q_N}(\boldsymbol{i})$ or NDH) with respect to the amount of ligand injected ($\boldsymbol{L_T}\ \Delta\boldsymbol{V_{inj}}(\boldsymbol{i})$)**,** as shown in Eqn (**10**),

$$\Delta Q_N(i) = \frac{\Delta Q(i)}{L_T\ \Delta V_{inj}(i)} \tag{10}$$

where, $\Delta\boldsymbol{V_{inj}}(\boldsymbol{i})$, is the volume of ligand injected. In NDH based ITC data analysis, we use, $\Delta\boldsymbol{Q_N}(\boldsymbol{i})$, the normalized area under the curve of peak corresponding to each injection as the dependent variable, and the total amount of protein $\boldsymbol{P_T}(\boldsymbol{i})$ and ligand $\boldsymbol{L_T}(\boldsymbol{i})$ as independent variables and fit it to the following Eqn (**11**),

$$\Delta Q_N(i) = \varDelta H\ (V_0[\ \Delta PL(i)]) \tag{11}$$

In the above model, $\Delta\boldsymbol{PL}(\boldsymbol{i})$ Eqn (**6**), in turn is dependent on $\boldsymbol{L}(\boldsymbol{i})$**,** Eqn (**7**), and constants, $\boldsymbol{P_T}(\boldsymbol{i})$ and ligand $\boldsymbol{L_T}(\boldsymbol{i})$.

#### 1.1.2. Time domain method

Following, the kinetic mechanism, Eqn (**1**), we can write the rate equations for $\boldsymbol{L}$, $\boldsymbol{P}$, and $\boldsymbol{PL}$ as,

$$\frac{d[L]}{dt} = -k_1[P][L] + k_{-1}[PL] \tag{12}$$

$$\frac{d[P]}{dt} = -k_1[P][L] + k_{-1}[PL] \tag{13}$$

$$\frac{d[PL]}{dt} = +k_1[P][L] - k_{-1}[PL] \tag{14}$$

Though $[\boldsymbol{L}]$, $[\boldsymbol{P}]$, and $[\boldsymbol{PL}]$ are not explicitly represented as $[\boldsymbol{L}(\boldsymbol{t})]$, $[\boldsymbol{P}(\boldsymbol{t})]$, and $[\boldsymbol{PL}(\boldsymbol{t})]$, it is implied that these variables are time dependent in the above model. In an ITC experiment, the ligand is injected in a discrete and sequential manner. After each injection ($\boldsymbol{i}$) the concentration of $[\boldsymbol{PL}]$ varies dynamically for a transient period until it reaches an equilibrium concentration $[\boldsymbol{PL}(\boldsymbol{i})]$. The change in the concentration of $[\boldsymbol{PL}]$ is accompanied by heat release or absorbed. This transient heat released or absorbed in the sample cell gives rise to a difference in temperature between the sample cell and the reference cell. Calorimeter measures this temperature difference in terms of 'differential power' (DP) or simply 'power' ($\boldsymbol{P_s}$; Cal/sec). The time profile of DP appears as an assymetrical gaussian peak in the thermogram. In thermodynamics, the DP or 'power' by definition is the rate of change of internal energy or heat ($\boldsymbol{dq}$) with respect to time ($\boldsymbol{dt}$), Eqn (**15**).

$$P_s(t(j)) = \frac{\boldsymbol{dq}(t(j))}{\boldsymbol{dt}} = \frac{q_{t(j)} - q_{t(j-1)}}{t(j) - t(j-1)} \tag{15}$$

In the above Eqn (**15**), $\boldsymbol{dq}(\boldsymbol{t}(\boldsymbol{j}))$ is the change in heat, $\boldsymbol{q}$, over the time interval, $\boldsymbol{t}(\boldsymbol{j}-\boldsymbol{1})$ to $\boldsymbol{t}(\boldsymbol{j})$. The term $\boldsymbol{dq}(\boldsymbol{t}(\boldsymbol{j}))$ in Eqn (**15**) is similar to that of the LHS term $\Delta\boldsymbol{Q}(\boldsymbol{i})$ in Eqn (**8**), expect that the former represents the instantaneous or very small heat change occuring between the time interval $\boldsymbol{t}(\boldsymbol{j}-\boldsymbol{1})$ and $\boldsymbol{t}(\boldsymbol{j})$, whereas, $\Delta\boldsymbol{Q}(\boldsymbol{i})$, represents the cumulative heat difference between the entire time interval that represents a complete injection, i.e. the time period, $\boldsymbol{t_0}$, starting from the end of the $(\boldsymbol{i-1})^{\boldsymbol{th}}$ injection till, $\boldsymbol{t_{end}}$, the end of $\boldsymbol{i^{th}}$ injection Eqn(**16**).

$$\int_{j=t_0}^{j=t_{end}} \frac{dq(j)}{dt} = Q|_{i-1}^{i} = Q_i - Q_{i-1} = \Delta Q(i) \tag{16}$$

Since we know that the rate of change of heat is proportional to the rate of change of complex formation we write Eqn (**17**),

$$\frac{dq}{dt} = \Delta H\, V_0 \frac{d[PL]}{dt} \tag{17}$$

Using Eqn (**15**) and Eqn (**17**), we can explicitly write the expression for $\boldsymbol{P_s}$ in terms of $[\boldsymbol{PL}]$ as Eqn (**18**),

$$P_s(t(j)) = \frac{dq}{dt} = \Delta H\, V_0 \frac{d[PL]}{dt} \tag{18}$$

Though, Eqn (**18**) is sufficient to simulate the thermogram, we preferred to use the following finite difference formula Eqn (**19**), in all our simulation and analysis mentioned in this work.

$$P_s(t(j)) = \Delta H\, V_0 \frac{\Delta[PL]}{\Delta t} = \Delta H\, V_0 \left(\frac{[PL(t(j))] - [PL(t(j-1))]}{t(j) - t(j-1)}\right) \tag{19}$$

Where, $[\boldsymbol{PL}(\boldsymbol{t}(\boldsymbol{j}-\mathbf{1}))]$ and $[\boldsymbol{PL}(\boldsymbol{t}(\boldsymbol{j}))]$ are concentrations of complex, $\boldsymbol{PL}$ at $\boldsymbol{t}(\boldsymbol{j}-\mathbf{1})$ and $\boldsymbol{t}(\boldsymbol{j})$, respectively. To avoid confusion in notation used here, we reserve '$\boldsymbol{i}$' as the index for injection and '$\boldsymbol{j}$', as the index for time points. The above Eqn (**19**) considers only the heat change due to binding process, if we take into account the heat of dilution and other non specific heat ($\boldsymbol{Q_{ns}}$), then we write Eqn (**20**),

$$\frac{dQ_T}{dt} = \frac{dQ_{Bind}}{dt} + \frac{dQ_{Dil}}{dt} + \frac{dQ_{ns}}{dt} \tag{20}$$

$$= \Delta H_{Bind}\, V_0 \frac{[PL(t(j))] - [PL(t(j-1))]}{t(j) - t(j-1)} + \Delta H_{Dil}\, V_0 \frac{[L(t(j))] - [L(t(j-1))]}{t(j) - t(j-1)} +$$

$$\Delta H_{ns}\, V_0 \frac{[Q_{ns}(t(j))]-[Q_{ns}(t(j-1))]}{t(j)-t(j-1)}$$

Where, $[\boldsymbol{L}(\boldsymbol{t}(\boldsymbol{j}-\mathbf{1}))]$ and $[\boldsymbol{L}(\boldsymbol{t}(\boldsymbol{j}))]$ are the concentrations of free ligand, $\boldsymbol{L}$ at $\boldsymbol{t}(\boldsymbol{j}-\mathbf{1})$ and $\boldsymbol{t}(\boldsymbol{j})$, respectively. In all the analysis carried out here, we have not considered contributions from non specific sources. The simulation of the ITC chromatogram for a two-state system without instrument response is presented in Figure 1 A,B. The parameters used for the simulation is summarized in Table SI 1.

*Table 1 Parameters used to simulate the thermograms of different models shown in Figure 1 (A,C,G,E) using dynamic approach.*

| **Model** | One site |
|---|---|
| **Parameters** | |
| Kinetic constants ($k$) | $1 \times 10^6$ $Hz$ ($k_1$)<br>1.0 ($Hz, k_{-1}$) |
| Stoichiometric constant | 1.0 ($m\ index$) |
| Thermodynamic constants ($\Delta H\ cals/mol$) | $-1 \times 10^4$ cals/mol ($\Delta H_{bind}$)<br>$-1 \times 10^2$ cals/mol ($\Delta H_{dil}$) |
| Injection, Cell parameters | 20 (Inj no)<br>4 µL(Inj vol)<br>230 µL (Cell vol) |
| Instrument response | 0.33 Hz ($k_{lig}$)<br>0.33 Hz ($k_{+h}, k_{-h}$) |
| Other parameters | 20 µM (Prot)<br>200 µM (Lig) |
| Integration parameters per peak in the thermogram | 0 - 100 s (each inj)<br>50 (data points) |

### 1. 2. Model with instrument response (Lumped model)

In this section, we outline the lumped modelling approach adopted by Dumas etal that incorporates the instrument response for (1) delay in mixing or dilution of the ligand after each injection, (2) delay in detection of heat released or absorbed by the detector. The instrument response is modelled here as a first order system independent to that of the binding mechanism. The modelling is carried out in a sequential manner as follows, (1) the time profile of the 'injected' ligand concentration ($\boldsymbol{L_{T_{inj}}(t)}$) is proposed according to an experimental design (2) the 'measured' Ligand ($\boldsymbol{L_{T_m}(t)}$) or Ligand available to bind to the protein is calculated from the 'injected' ligand, $\boldsymbol{L_{T_{inj}}(t)}$, considering first order instrument response (3) A suitable kinetic mechanism is proposed to explain the binding mechanism and the concentrations of $\boldsymbol{L(t)}$, $\boldsymbol{P(t)}$, and $\boldsymbol{PL(t)}$ are calculated. (4) The 'ideal' power, $\boldsymbol{\overline{P}_s(t)}$ required to simulate the ITC time profile is calculated. (5) The 'measured', $\boldsymbol{\overline{P}_m(t)}$, which represents the thermogram is calculated from the ideal, $\boldsymbol{\overline{P}_s(t)}$ by considering first order intrument response (6) Each peak in the thermogram corresponding to individual injections are integrated and normalized with respect to the injected ligand ($\Delta\boldsymbol{L_{T_{inj}}(i)}$) and represented as NDH data.

#### 1.2.1. Injected ligand profile

The time profile of the ligand injected into the cell can be either linear, or non-linear (convex or concave) depending on the experimental design. For example, a linear model would be, $\boldsymbol{L_{T_{inj}}(t) = k_{inj} * t}$, where, $\boldsymbol{k_{inj}}$ is the slope or the rate at which the ligand is injected. If, $\boldsymbol{\Delta V}$, is the volume of ligand injected over a period of time period, $\boldsymbol{t_0}$ to $\boldsymbol{t_{end}}$, then, $\boldsymbol{k_{inj}} = \frac{\boldsymbol{\Delta V}}{\boldsymbol{t_{end} - t_0}}$.

#### 1.2.2. Measured ligand profile

The measured ligand ($\boldsymbol{L_{T_m}(t)}$) profile is calculated from the injected ligand profile ($\boldsymbol{L_{T_{inj}}(t)}$) by solving the following first order differential equation,

$$\frac{dL_{T_m}}{dt} = \frac{1}{\tau_{lig}}\left(L_{T_{inj}}(t) - L_{T_m}(t)\right) \qquad (21)$$

In the above equation, $\boldsymbol{\tau_{lig}}$, is the delay or the instrument response time, required by the system to reach an equilibrium concentration from the time of injection. If the ligand mixing/stirring is efficient and fast, then the 'instrument response time', $\boldsymbol{\tau_{lig}}$ would be closer to zero.

#### 1.2.3. Binding mechanism

Assuming a binding mechanism, such as the single site binding Eqn (**1**), the time profiles of the free ligand $\boldsymbol{L(t)}$, free protein, $\boldsymbol{P(t)}$, and bound protein or complex $\boldsymbol{PL(t)}$ can be easily calculated by numerically solving the set of coupled differential equations mentioned earlier Eqns (**13**, **14**, **15**). While solving the ODE's, the initial condition of the first time point after the injection is as follows, Eqns (**22**, **23**, **24**),

$$[L(t(0))]_0 = L_{T_m}(0) \tag{22}$$

$$[P(t(0))]_0 = P_T(0) \tag{23}$$

$$[PL(t(0))]_0 = 0 \tag{24}$$

Where, $\boldsymbol{L_{T_m}(0)}$, $\boldsymbol{P_T(0)}$ are the intial concentration of the total ligand and total protein present in the cell, respectively, prior to the injection. The initial condition for the second and subsequent time points are given by, Eqns (**25**, **26**, **27**),

$$[L(t(j))]_0 = \left[L_{T_m}(t(j))\right] + \ [L(t(j-1))] \tag{25}$$

$$[P(t(j))]_0 = \ [P(t(j-1))] * DF \tag{26}$$

$$[PL(t(j))]_0 = [PL(t(j-1))] \tag{27}$$

Where, $\boldsymbol{L_{T_m}}(\boldsymbol{t(j)})$, is the concentration of the measured total ligand obtained by solving Eqn (**21**). DF is the dilution factor considered for correcting the additional volume of ligand injected in to the cell and the excess volume of solution overflow out of the cell into the stem region.

The effect of dilution on concentration of protein and ligand is considered here. At the outset, the initial concentration of the protein is $\boldsymbol{P_{T_0}}$, and the cell volume is $\boldsymbol{V_0}$. The total amount of protein present is given by, $\boldsymbol{P_{T_0} V_0}$. If a small volume of ligand $\Delta \boldsymbol{V_1}$, is added to this solution, an equal volume of the protein inside the cell is displaced into the stem region. So the apparent change in concentration due to this instantaneous displacement is given by Eqn (**28**),

$$P_{T_1} = \frac{\underbrace{P_{T_0} V_0}_{protein\ in\ cell} - \underbrace{P_{T_0} \Delta V_1}_{protein\ displaced\ to\ the\ stem}}{V_0} = P_{T_0}\left(\frac{V_0 - \Delta V_1}{V_0}\right) \tag{28}$$

Similarly, the second injection volume of $\Delta \boldsymbol{V_2}$, will dilute the protein to an apparent concentration as follows Eqn (**29**),

$$P_{T_2} = P_{T_1}\left(\frac{V_0 - \Delta V_2}{V_0}\right) = P_{T_0}\left(\frac{V_0 - \Delta V_1}{V_0}\right)\left(\frac{V_0 - \Delta V_2}{V_0}\right) \tag{29}$$

After $\boldsymbol{n}$ th injection the apparent concentration of protein can be written as Eqn (**30**),

$$P_{T_n} = P_{T_0}\left(\prod_{i=1}^{n}\left(\frac{V_0 - \Delta V_i}{V_0}\right)\right) = P_{T_0}\left(\prod_{i=1}^{n}\left(1 - \frac{\Delta V_i}{V_0}\right)\right) \tag{30}$$

Similarly, for the ligand we consider the initial concentration of ligand in the cell to be $\boldsymbol{L_{T_0}}$, and the cell volume is $\boldsymbol{V_0}$**.** If we add a small volume of ligand $\Delta \boldsymbol{V_1}$ of concentration $\boldsymbol{L_{T_{inj}}}$, then the apparent change in concentration of ligand inside the cell due to addition and dilution effect is given by Eqn (**31**),

$$L_{T_1} = \frac{\underbrace{L_{T_0}V_0}_{inside\ cell} - \underbrace{L_{T_0}\Delta V_1}_{displaced\ to\ stem} + \underbrace{L_{T_{inj}}\Delta V_1}_{injected\ into\ cell}}{V_0} = L_{T_{inj}}\left(\frac{\Delta V_1}{V_0}\right) \tag{31}$$

Since, $\boldsymbol{L_{T_0} = 0}$, we have only the injected ligand contributing to the ligand concentration inside the cell for the first injection. The subsequent injection of $\boldsymbol{\Delta V_1}$, results in Eqn (**32**),

$$L_{T_2} = L_{T_1}\left(\frac{V_0 - \Delta V_2}{V_0}\right) + L_{T_{inj}}\left(\frac{\Delta V_2}{V_0}\right) \tag{32}$$

After $\boldsymbol{n}$ th injection the apparent concentration of ligand inside the cell can be written as Eqn (**33**),

$$L_{T_n} = L_{T_{(n-1)}}\left(\frac{V_0 - \Delta V_n}{V_0}\right) + L_{T_{inj}}\left(\frac{\Delta V_n}{V_0}\right) \tag{33}$$

Successive substitutions of previous concentrations leads to the following Eqn (**34**),

$$L_{T_n} = L_{T_{inj}}\left(1 - \left(\prod_{i=1}^{n}\left(1 - \frac{\Delta V_i}{V_0}\right)\right)\right) \tag{34}$$

#### 1.2.4. Ideal power profile

The ITC chromatogram is represented in terms of 'power', $\boldsymbol{\bar{P}_s(t(j))}$, or the rate of change of heat $\boldsymbol{q_s(t(j))}$, as follows Eqn (**35**),

$$\begin{aligned}\bar{P}_s(t(j)) &= \frac{dq_s(t(j))}{dt} = \Delta H\ V_0\ \frac{d[\,PL(t(j))]}{dt} \\ &\equiv \Delta H\ V_0\ \frac{([PL(t(j))] - [PL(t(j-1))])}{(t(j) - t(j-1))}\end{aligned} \tag{35}$$

The above expression for power is considered to be ideal because the instrument response is not yet taken into account.

#### 1.2.5. Measured power profile

The measured power ($\boldsymbol{\bar{P}_m(t(j))}$) is calculated from the ideal power ($\boldsymbol{\bar{P}_s(t(j))}$) by solving the following first order differential equation,

$$\frac{d\bar{P}_m(t(j))}{dt} = \frac{1}{\tau_{heat}}\left(\bar{P}_s(t(j)) - \bar{P}_m(t(j))\right) \quad (36)$$

The experimentally measured ITC chromatogram is nothing but the time profile of the measured 'power', $\boldsymbol{\bar{P}_m(t(j))}$ .

#### 1. 3. Model with instrument response (Kinetic sequential model)

In this section, we outline the kinetic based modelling approach that is identical to that of the lumped modelling approach. As before, both the instrument response in (1) ligand dilution and (2) heat detection, are modelled here as first order system and independent of the binding kinetics.

The 'measured ligand' is modelled based on the 'injected ligand' using the following first order kinetic model,

$$L_{T_{inj'}} \xrightarrow{k_{lig}} L_{T_m} \quad (37)$$

Based on which the time profile of the 'measured ligand' can be calculated using the following rate equation,

$$\frac{dL_{T_m}(t)}{dt} = k_{lig}L_{T_{inj'}}(t) = k_{lig}\left(L_T(t) - L_{T_m}(t)\right) \quad (38)$$

Where, $\boldsymbol{L_T(t)}$, is the total ligand concentration inclusive of both injected and measured ligand ($\boldsymbol{L_T(t) = L_{T_{inj'}}(t) + L_{T_m}(t)}$). The term $\boldsymbol{L_{T_{inj'}}(t)}$ represents the unavailable part of the total injected ligand which is yet to be dispersed into the solution. This term, $\boldsymbol{L_{T_{inj'}}(t)}$, is not to be confused with the $\boldsymbol{L_{T_{inj}}(t)}$, mentioned in Eqn (**21**), which represented the total amount of ligand injected which included both the dispersed and the undispersed ligand.

$\boldsymbol{L_{T_m}(t)}$, as calculated above is used to get the time profiles of the free ligand $\boldsymbol{L(t)}$, free protein, $\boldsymbol{P(t)}$, and bound protein or complex $\boldsymbol{PL(t)}$ based on an assumed binding mechanism. The ideal power is calculated in a similar way as explained in Sec 1.2.4. Given the ideal power, the measured power can be modelled using the following kinetics Eqn (**39**),

$$\bar{P}_s \xrightarrow{k_{heat}} \bar{P}_m \tag{39}$$

The rate equation used to determine the time profile of the ideal power can be written as, Eqn (**40**)

$$\frac{d\bar{P}_m(t)}{dt} = k_{heat}\bar{P}_s(t) = k_{heat}(\bar{P}_T(t) - \bar{P}_m(t)) \tag{40}$$

Where, $\boldsymbol{\bar{P}_T(t)}$, is the total power inclusive of both the ideal and measured heat ($\boldsymbol{\bar{P}_T(t) = \bar{P}_s(t) + \bar{P}_m(t)}$). When numerically integrating Eqn (**40**), the term $\boldsymbol{k_{heat}\bar{P}_T(t)}$ will be provided as the initial condition and is nothing but $\boldsymbol{\bar{P}_s(0)}$, which is obtained through Eqn (**35**). The equivalences of lumped model and the kinetic model are explained in detail in SI Sec 1. 5.

#### 1. 4. Model with instrument response (Kinetic parallel model)

In this section, we outline the kinetic based modelling approach, which incorporates instrument response within the binding mechanism. This approach allows for the simultaneous modelling of instrument responses and binding mechanism with in a simple kinetic framework. The kinetic mechanism can be proposed as follows,

$$L_{T_{inj'}} \xrightarrow{k_{lig}} L \tag{41}$$

Eqn (41) accounts for the instrument response of the ligand dispersion.

$$P + L \underset{k_{-1}}{\overset{k_1}{\rightleftharpoons}} PL \tag{42}$$

Eqn (42) accounts for the binding mechanism of the available ligand with the protein

$$PL \underset{k_{-h}}{\overset{k_{+h}}{\rightleftharpoons}} \overline{PL} \tag{43}$$

Eqn (43) accounts for the instrument response of the heat released or absorbed due to binding process by making the ‘power’ proportional to rate of change of $\overline{PL}$, rather than $PL$ itself.

$$L \underset{k_{-h}}{\overset{k_{+h}}{\rightleftharpoons}} \bar{L} \tag{44}$$

Eqn (44) accounts for the instrument response associated with heat of ligand dilution, by making the ‘power’ proportional to the rate of change of $\bar{L}$, rather than $L$ itself.

The rate equation for the above kinetic mechanism can be written as, Eqns (45 - 50),

$$\frac{d\left[L_{T_{inj'}}\right]}{dt} = -k_{lig}\left[L_{T_{inj'}}\right] \tag{45}$$

$$\frac{d[L]}{dt} = k_{lig}\left[L_{T_{inj'}}\right] - k_1[P][L] + k_{-1}[PL] \tag{46}$$

$$\frac{d[P]}{dt} = -k_1[P][L] + k_{-1}[PL] \tag{47}$$

$$\frac{d[PL]}{dt} = +k_1[P][L] - k_{-1}[PL] \tag{48}$$

$$\frac{d[\overline{PL}]}{dt} = +k_{+h}[PL] - k_{-h}[\overline{PL}] \quad (49)$$

$$\frac{d[\bar{L}]}{dt} = +k_{+h}[L] - k_{-h}[\bar{L}] \quad (50)$$

The above set of differential equations can be solved simultaneously to obtain the ITC time profile. Fig (1 A-H) shows, the simulated ITC chromatogram using all the four approaches explained in the sections (1.1 to 1.4).

#### 1. 5. Modelling the instrument response of Ligand dilution and 'Power' measurement

The time lag between the availability of the ligand ($L_{T_m}$) after injection ($L_{T_{inj}'}$) and the calculation of 'measured power', ($\bar{P}_m(t)$) from the 'ideal power', ($\bar{P}_s(t)$), can be modelled using several approaches. (1) Convolution based appoach, (2) Lumped modelling, (3) Kinetic approach. In this section, we derive and show the equivalence of all these approaches for the instrument response considerd in 'measured power' and can be easily extended to model instrument response of 'measured ligand' concentration.

##### 1.5.1. Convolution based modelling appoach

In this approach the 'ideal power', ($\boldsymbol{\bar{P}_s(t)}$), is convoluted with an kernel function, ($\boldsymbol{\bar{K}}$) to model the 'measured power', ($\boldsymbol{\bar{P}_m(t)}$), Eqn (**51**),

$$\bar{P}_m(t) = \bar{P}_s(t) \otimes \bar{K} \quad (51)$$

For a first order system, the kernel function, ($\bar{\boldsymbol{K}}$) is modelled as a mono exponential decay function with a time lag constant $\boldsymbol{\tau_{heat}}$, Eqn (**52**),

$$\bar{K} = \frac{1}{\tau_{heat}} \, e^{-\left(\frac{t}{\tau_{heat}}\right)} \tag{52}$$

Substituting, Eqn (**52**) into Eqn (**51**) yields,

$$\bar{P}_m(t) = \bar{P}_s(t) \otimes \frac{1}{\tau_{heat}} \, e^{-\left(\frac{t}{\tau_{heat}}\right)} \tag{53}$$

Expanding the above Eqn (**53**) based on the definition of convolution, $\boldsymbol{f(t) \otimes g(t) = \int_0^t f(\theta).\, g(t-\theta)\, d\theta}$, we have Eqn (**54**),

$$\bar{P}_m(t) = \int_0^t \bar{P}_s(\theta).\frac{1}{\tau_{heat}} \, e^{-\left(\frac{t-\theta}{\tau_{heat}}\right)} \, d\theta \tag{54}$$

Differentiating the above Eqn (**54**) with respect to $\boldsymbol{t}$, yeilds, Eqn (**55**),

$$\bar{P}_m{}'(t) = \frac{d}{dt}\left(\int_0^t \bar{P}_s(\theta).\frac{1}{\tau_{heat}} \, e^{-\left(\frac{t-\theta}{\tau_{heat}}\right)} \, d\theta\right) \tag{55}$$

For the term within the inner integral, both $\boldsymbol{t}$ and $\boldsymbol{\tau_{heat}}$ are constants with respect to θ, and hence can be taken out of the integral for simplification Eqn (**56**).

$$\bar{P}_m{}'(t) = \frac{d}{dt}\left(\left(\frac{1}{\tau_{heat}} \, e^{-\left(\frac{t}{\tau_{heat}}\right)}\right)\left(\int_0^t \bar{P}_s(\theta).\; e^{\left(\frac{\theta}{\tau_{heat}}\right)} \, d\theta\right)\right) \tag{56}$$

Differentiating the Eqn(**56**) using chain rule yeilds, Eqn (**57**)

$$\bar{P}_m{}'(t) = \left(\frac{1}{\tau_{heat}}\, e^{-\left(\frac{t}{\tau_{heat}}\right)}\right) \underbrace{\frac{d}{dt}\left(\left(\int_0^t \bar{P}_s(\theta).\, e^{\left(\frac{\theta}{\tau_{heat}}\right)}\, d\theta\right)\right)}_{A} + \left(\int_0^t \bar{P}_s(\theta).\, e^{\left(\frac{\theta}{\tau_{heat}}\right)}\, d\theta\right)\frac{d}{dt}\left(\frac{1}{\tau_{heat}}\, e^{-\left(\frac{t}{\tau_{heat}}\right)}\right) \quad (57)$$

In the above Eqn (**57**), to simplify the term $\boldsymbol{A}$, we use the fundamental theorem of the differentiation of an integral function, Eqn (**58**)

$$\frac{d}{dt}\left(\int_0^t f(x)\, dx\right) = f(t) \quad (58)$$

Notably, in the inner integral $\boldsymbol{t}$, appears as a constant (upper limit for $\boldsymbol{x}$), whereas, in the outer differentiation, t is considered to be a variable. The differentiation of such an integral simply becomes $\boldsymbol{f}(\boldsymbol{x}) \rightarrow \boldsymbol{f}(\boldsymbol{t})$.

$$\bar{P}_m{}'(t) = \left(\frac{1}{\tau_{heat}}\, e^{-\left(\frac{t}{\tau_{heat}}\right)}\right)\left(\bar{P}_s(t).\, e^{\left(\frac{t}{\tau_{heat}}\right)}\right) + \left(\int_0^t \bar{P}_s(\theta).\, e^{\left(\frac{\theta}{\tau_{heat}}\right)}\, d\theta\right)\underbrace{\frac{d}{dt}\left(\frac{1}{\tau_{heat}}\, e^{-\left(\frac{t}{\tau_{heat}}\right)}\right)}_{B} \quad (59)$$

Differentiating term $\boldsymbol{B}$ of Eqn (**59**) yeilds Eqn (**60**),

$$\bar{P}_m{}'(t) = \left(\frac{1}{\tau_{heat}}\, e^{-\left(\frac{t}{\tau_{heat}}\right)}\right)\left(\bar{P}_s(t).\, e^{\left(\frac{t}{\tau_{heat}}\right)}\right) + \underbrace{\left(\int_0^t \bar{P}_s(\theta).\, e^{\left(\frac{\theta}{\tau_{heat}}\right)}\, d\theta\right)}_{C}\left(\frac{1}{\tau_{heat}}\right)\underbrace{e^{-\left(\frac{t}{\tau_{heat}}\right)}\left(-\frac{1}{\tau_{heat}}\right)}_{D} \quad (60)$$

Rearranging the terms in $\boldsymbol{C}$ and bring part of the terms in $\boldsymbol{D}$,inside the integral yeilds Eqn (**61**),

$$\bar{P}_m{}'(t) = \left(\frac{1}{\tau_{heat}}\, e^{-\left(\frac{t}{\tau_{heat}}\right)}\right)\left(\bar{P}_s(t).\, e^{\left(\frac{t}{\tau_{heat}}\right)}\right) + \left(-\frac{1}{\tau_{heat}}\right)\left(\int_0^t \bar{P}_s(\theta).\, e^{\left(\frac{\theta}{\tau_{heat}}\right)}\, e^{-\left(\frac{t}{\tau_{heat}}\right)}\left(\frac{1}{\tau_{heat}}\right) d\theta\right) \tag{61}$$

Rearranging the above Eqn (**62**) yeilds,

$$\bar{P}_m{}'(t) = \left(\frac{1}{\tau_{heat}}\, e^{-\left(\frac{t}{\tau_{heat}}\right)}\right)\left(\bar{P}_s(t).\, e^{\left(\frac{t}{\tau_{heat}}\right)}\right) + \left(-\frac{1}{\tau_{heat}}\right)\left(\underbrace{\int_0^t \bar{P}_s(\theta).\frac{1}{\tau_{heat}}\, e^{-\left(\frac{t-\theta}{\tau_{heat}}\right)} d\theta}_{E}\right) \tag{62}$$

The product of the exponent parts of the first term in RHS reduces to 1, and the second term , $\boldsymbol{E}$ , in RHS is nothing but $\boldsymbol{\bar{P}_m(t)}$, hence, the Eqn (**62**) on simplification yeilds Eqn (**63**),

$$\bar{P}_m{}'(t) = \frac{1}{\tau_{heat}}\left(\bar{P}_s(t) - \bar{P}_m(t)\right) \tag{63}$$

#### 1.5.2. Lumped modelling appoach

Often engineers tend to model their system using fundamental electrical/electronic components such as resistors, capacitors and inductors. Equivalence of electrical and mechanical components based modelling are well established. Combination of these basis elements in specific order is often used in modelling the system as accurate as possible, so as to study the time domain and frequency domain responses. Here, we use two simple cases, $\boldsymbol{\bar{R}\bar{C}}$ and $\boldsymbol{\bar{R}\bar{L}}$ systems which can be used to model the first order instrument response of the 'measured power'. In such modelling approach, two variables are defined namely, 'through variable' and 'across variable'. Since current '$\boldsymbol{I}$' passes through the system it is defined to be the 'through variable' and since, the voltage '$\boldsymbol{V}$' is measured across two points in a circuit, it is defined to be 'across variable' [1]. Extensive use of

'Kirchoff's first and second law enable us to obtain differential equations relating both these variables.

##### 1.5.2.1. RL system

Firstly, we consider a system with resistor '$\boldsymbol{\bar{R}}$' and inductor '$\boldsymbol{\bar{L}}$', connected serially. The voltage across resistor, $\boldsymbol{V_R}$ , is linearly proportional to current , $\boldsymbol{I}$, and is modelled as Eqn (**64**),

$$V_R = R\,I \tag{64}$$

Where, $\boldsymbol{R}$, is the resistance constant. The voltage across inductor , $\boldsymbol{V_L}$ , is the derivative of current and is modelled as, Eqn (**65**),

$$V_L = L\,\frac{dI}{dt} \tag{65}$$

Where, $\boldsymbol{L}$, is the inductance constant.The total voltage across both the resistor and inductor is given by, Eqn (**66**),

$$V = V_R + V_L = \; R\,I + L\,\frac{dI}{dt} \tag{66}$$

If we represent**,** $\boldsymbol{\bar{P}_m(t) = R\,I,}$ then its derivative is**,** $\boldsymbol{\bar{P}_m{}'(t) = R\,I';}$ similarly, we represent $\boldsymbol{\bar{P}_s(t) = V}$. Substituting these new variable into the Eqn (**66**) yeilds Eqn (**67**),

$$\bar{P}_s(t) = \; \bar{P}_m(t) + \tfrac{L}{R}\,\bar{P}_m{}'(t) \tag{67}$$

Rearrangement of the above Eqn (**67**) yeilds, Eqn (**68**),

$$\bar{P}_m{}'(t) = \frac{R}{L}\,\left(\bar{P}_s(t) - \bar{P}_m(t)\right) \tag{68}$$

If we represent $\frac{1}{\boldsymbol{\tau_{heat}}} = \frac{\boldsymbol{R}}{\boldsymbol{L}}$, then we obtain the model Eqn (**68**), identical to Eqn (**36**) obtained in previous section.

#### 1.5.2.2. RC system

Here, we consider a system with resistor '$\overline{\boldsymbol{R}}$' and capacitor '$\overline{\boldsymbol{C}}$', connected serially. The voltage across capacitor, $\boldsymbol{V_C}$ , is an integral of current and is modelled as, Eqn (**69**),

$$V_C = \frac{1}{C} \int I \ dt \tag{69}$$

Where, $\boldsymbol{C}$, is the capacitance constant. The total voltage across both the resistor and capacitor is given by Eqn (**70**),

$$V = V_R + V_C = \ R\,I + \frac{1}{C} \int I \ dt \tag{70}$$

Differentiating the above Eqn (**70**), leads to Eqn (**71**),

$$V' = \ R\,I' + \frac{1}{C}\,I \tag{71}$$

If we represent, $\overline{\boldsymbol{P}}_{\boldsymbol{m}}(\boldsymbol{t}) = \frac{\boldsymbol{1}}{\boldsymbol{C}}\,\boldsymbol{I}$, then its derivative is, $\overline{\boldsymbol{P}}_{\boldsymbol{m}}{}'(\boldsymbol{t}) = \frac{\boldsymbol{1}}{\boldsymbol{C}}\,\boldsymbol{I'}$**;** similarly, we represent $\overline{\boldsymbol{P}}_{\boldsymbol{s}}(\boldsymbol{t}) = \boldsymbol{V'}$. Substituting these new variable into the Eqn (**71**) yeilds Eqn (**72**),

$$\bar{P}_s(t) = RC\,\bar{P}_m{}'(t) + \ \bar{P}_m(t) \tag{72}$$

Rearrangement of the above Eqn (**72**) yeilds, Eqn (**73**)

$$\bar{P}_m{}'(t) = \frac{1}{RC}\left(\bar{P}_s(t) - \bar{P}_m(t)\right) \tag{73}$$

If we represent $\frac{\mathbf{1}}{\boldsymbol{\tau_{heat}}} = \frac{\mathbf{1}}{\boldsymbol{RC}}$, then we obtain the model Eqn (**73**), identical to Eqn (**36**) obtained in previous section.

The comparison of Eqn (68) and Eqn (73) suggests that the conserved quantity, the electric potential energy ($V$) or its derivative ($V'$), represents the 'ideal power', ($\bar{P}_s(t)$), where as the 'current' ($I$), represents the dissipated 'measured power', ($\bar{P}_m(t)$) .

#### 1.5.3. Kinetic modelling appoach

In kinetics based modelling we first propose a mechanism of how the reactants react together to form the product. Let's assume, a simple first order kinetics where a reactant, $\boldsymbol{L_1}$, yeilds a product, $\boldsymbol{L_2}$, Eqn (**74**),

$$L_1 \overset{k_1}{\rightarrow} L_2 \tag{74}$$

if we frame the rate equation for the product, $\boldsymbol{L_2}$ , we obtain, Eqn (**75**),

$$\frac{d[L_2]}{dt} = [L_2]' = k_1[L_1] \tag{75}$$

Rearranging the above equation to express, $L_1$, interms of, $L_2$, we get, Eqn (76),

$$[L_1] = \frac{[L_2]'}{k_1} \tag{76}$$

Using the mass balance for the species, $L$, the total concentration $L_T$, is represented as the sum of the reactant and product, Eqn (77),

$$L_T = L_1 + L_2 \tag{77}$$

Substituting Eqn (76), into Eqn (77), yeilds Eqn (78),

$$L_T = \frac{[L_2]'}{k_1} + L_2 \tag{78}$$

Rearranging the above equation yeilds, Eqn (79),

$$[L_2]' = k_1 \, (L_T - L_2) \tag{79}$$

If we subtitute $\boldsymbol{L_2 = \bar{P}_m(t)}$ , $\boldsymbol{L_T = \bar{P}_s(t)}$, and, $\boldsymbol{k_1 = \frac{1}{\tau_1}}$ we obtain the the model Eqn (**79**), identical to that of the Eqn (**36**) obtained in previous section.

The comparison of Eqn (68) and Eqn (79) suggests that the conserved quantity, the total mass ($L_T$) represents the 'ideal power', where as the final product ($L_2$), represents the dissipated 'measured power'.

The kinetic modelling enable us to easily model the higher order instrument response, e.g. second order, as follows, Eqn (80),

$$L_1 \overset{k_1}{\rightarrow} L_2 \overset{k_2}{\rightarrow} L_3 \tag{80}$$

Here, the rate equation of $L_3$, and , $L_2$ , Eqn (81, 82),

$$\frac{d[L_3]}{dt} = [L_3]' = k_2[L_2] \tag{81}$$

$$\frac{d[L_2]}{dt} = [L_2]' = k_1[L_1] \tag{82}$$

From Eqn (81), we obtain, Eqn (83),

$$[L_2] = \frac{[L_3]'}{k_2} \tag{83}$$

From Eqn (82), we obtain, Eqn (84),

$$[L_1] = \frac{[L_2]'}{k_1} \tag{84}$$

We derivate Eqn (83), to get Eqn (85),

$$[L_2]' = \frac{[L_3]''}{k_2} \tag{85}$$

Substituting Eqn (85) into Eqn (84), yeilds, Eqn (86),

$$[L_1] = \frac{[L_3]''}{k_1\, k_2} \tag{86}$$

The mass balance for the species , $L$, is given by, $L_T$, Eqn (87),

$$L_T = L_1 + L_2 + L_3 \tag{87}$$

Substituting Eqn (86, 83), into Eqn (87), yeilds, Eqn (88),

$$L_T = \frac{[L_3]''}{k_1\, k_2} + \frac{[L_3]'}{k_2} + L_3 \tag{88}$$

Rearranging the Eqn (88) yeilds, Eqn (89),

$$[L_3]'' + k_1[L_3]' + k_1 k_2 (L_3 - L_T) = 0 \tag{89}$$

If we substitute, $L_3 = \bar{P}_m(t), L_T = \bar{P}_s(t),\ k_1 = \frac{1}{\tau_1}$ , and, $k_2 = \frac{1}{\tau_2}$, into Eqn (89), we obtain the model for second order instrument response, Eqn (90),

$$\bar{P}_m(t)'' + \frac{1}{\tau_1}\bar{P}_m(t)' + \frac{1}{\tau_1 \tau_2}(\bar{P}_m(t) - \bar{P}_s(t)) = 0 \tag{90}$$

### 2. Simulation of SPR profile of a single site binding mechanism

SPR comprises of three distinct phases namely, association, dissociation and regeneration. The protein or the macromolecular species is covalently attached to the surface of the SPR sensor, whereas, the ligand is continuously equilibriated over the sensor through an isocratic flow. The kinetic mechanism can be proposed as follows, Eqn (147),

$$L_{T_{in}} \underset{k_{-lig}}{\overset{k_{lig}}{\rightleftharpoons}} L \tag{91}$$

Where, $L_{T_{in}}$, is the concentration of the ligand constantly flowing through the SPR system during the association phase and $L$, is the actual concentration of the ligand available for binding. $k_{lig}$, $k_{-lig}$ are the forward and reverse rates that determines the ligand ($L_{T_{in}}$) flows through the sensor is available in the form of $L$ for further binding kinetics. Eqn (147) is in a form that accounts for the instrument response due to ligand dilution or its availability to bind to protein. Altogeather, Eqn (147), accounts for the instrument response of the ligand dilution from the center of the 'cell' into bulk of the solution.

$$P + L \underset{k_{-1}}{\overset{k_1}{\rightleftharpoons}} PL \tag{92}$$

Where, $k_1$, $k_{-1}$, are the forward and reverse kinetic rate constants of the complex formation, respectively. Eqn (148) accounts for the binding mechanism of the available ligand, $L$ , with the protein, $P$, to form the complex, $PL$.

$$PL \underset{k_{-res}}{\overset{k_{+res}}{\rightleftharpoons}} \overline{PL} \tag{93}$$

Eqn (149) accounts for the instrument response related to the SPR's detector which is responsive to change in refractive index due to binding process. Here, the repsonse factor ($k_{+res}$) is related to the time delay $\left(\tau_{+res} = \frac{1}{k_{+res}}\right)$, observed between the instance of binding and the instance of signal detected by the sensor/detector. As a simplification, we will always assume that, ($k_{+res} = k_{-res}$), for the following discussions. The rate equations for the above kinetic mechanism can be written as Eqns (150-154),

$$\frac{d[L_{T_{in}}]}{dt} = -k_{lig}[L_{T_{in}}] + k_{lig}[L] \quad (94)$$

$$\frac{d[L]}{dt} = +k_{lig}[L_{T_{in}}] - k_{lig}[L] - k_1[P][L] + k_{-1}[PL] \quad (95)$$

$$\frac{d[P]}{dt} = -k_1[P][L] + k_{-1}[PL] \quad (96)$$

$$\frac{d[PL]}{dt} = +k_1[P][L] - k_{-1}[PL] \quad (97)$$

$$\frac{d[\overline{PL}]}{dt} = +k_{+res}[PL] - k_{-res}[\overline{PL}] \quad (98)$$

The above set of differential equations, Eqns (150-154), can be solved simultaneously to obtain, $[L_{T_{in}}]$, $[L]$, $[P]$, $[PL]$,$[\overline{PL}]$, at different time points. The initial condition or concentration of $L_{T_{in}}$, is kept constant through out the association regime; and is kept '0' during the dissociative and regeneration regimes. If we ought to consider a leak factor during the dissociation phase, which is also related to the residual baseline, then the concentration of $L_{T_{in}}$ can be modelled as a sum of constant part and an exponentially decaying function, $L_{T_{in}}(t) = L_{T_{in}}(0) + L_{T_{in}}(0)e^{-k_{leak}t}$; where, $k_{leak}$, is the leakage rate.

The detected response is always represented in terms of 'Response Unit', (RU), in the form of a sensogram. In Eqn (154), we have equated the 'RU' to be proportional to rate of change of $\overline{PL}$, rather than $PL$ itself, so as to account for the instrument response due to sensor. Given the time profile of $[\overline{PL}]$, the time profile of the 'sensogram', $RU(t(j))$, can be calculated as follows Eqn (155),

$$RU(t(j)) = G.\overline{PL}(t(j)) \tag{99}$$

Where, $G$, is the gain factor, (a constant); $\overline{PL}\ (t(j))$ is the concentration of the complex, $[\overline{PL}]$ at time $t(j)$.

*Table 2 Parameters used to simulate the sensograms of a single site binding using dynamic approach for cases with and without leakage.*

| | SPR |
|---|---|
| **Model** | Single site binding mechanism |
| **Parameters** | |
| Kinetic rate constants ($k$) | $1.0 \times 10^3$ Hz ($k_1$)<br>$1.0 \times 10^5$ Hz ($k_{-1}$) |
| Stoichiometric constant | 1 ($m\ index$) |
| Experimental time | 20s (Total)<br>10s (Association)<br>10s (Dissociation) |
| Instrument response | 1.0 ($k_{lig}$)<br>1.0 ($k_{+h}, k_{-h}$) |
| Other parameters | 10 μM (Prot)<br>1, 5, 7, 10, 15, 20, 25, 30 μM (Lig) |
| Integration parameters for sensogram | 20 s (Total expt)<br>100 (data points) |
| Leakage factor | 20% (L(0) %; Basal value in % to that of the Ligand conc flow in the sensor) |

| | |
|---|---|
| | $1.0 \times 10^8$ Hz ($k_{leak}$; Leak rate) |

#### 3. Sensitivity analysis for the fit model parameters

There are several methods available to assess the uncertainty of the optimized parameters obtained by fitting an experimental data to a specified model. Consider a model, where the output variable, $Y$, is dependent on the independent variable, $X$, and the constants, $x_1, x_2, \cdots, x_n$, as follows,

$$Y = g(X, x_1, x_2, \cdots, x_n) \tag{100}$$

If we optimize the parameters of the model, $x_1, x_2, \cdots, x_n$, for a given experimental data, the uncertainity in the fit parameters is often used to assess the sensitivity of the parameters for the given model.

In commonly used approaches such as derivative and variance based methods, the final fit parameters are varied slightly from its optimized value and the variance in the output values ($Y$) are back calculated. If the resultant output variance is minimal, then it implies that the uncertainty in the fit parameter is also minimal and is reliable. The underlying assumption in these methods is that the statistical moment, the variance, is correlated and proportional to uncertainty. Further, the fit parameters are assumed to be sampled from a "probabilistic" model distribution that resembles Gaussian. On the other hand, there are probability model independent approaches available to assess the uncertainty in the fit parameters E.g. density based approaches. In this approach, firstly, a probability density function (PDF) of the output variable ($Y$) is generated by randomly varying all the fit parameters. Because all the variables are varied it can be considered as an 'unconditional PDF'. Secondly, another PDF of the output ($Y$) is generated by varying all the parameters except the parameter of interest whose uncertainty has to be assessed. This PDF is commensurate with the 'conditional PDF'. The difference of divergence between the conditional and the unconditional PDF is a quantitative measure of the global sensitivity of that parameter. Finally, Kolmogorov Smirnov statistics is applied to assess the statistical significance of the obtained sensitivity values.

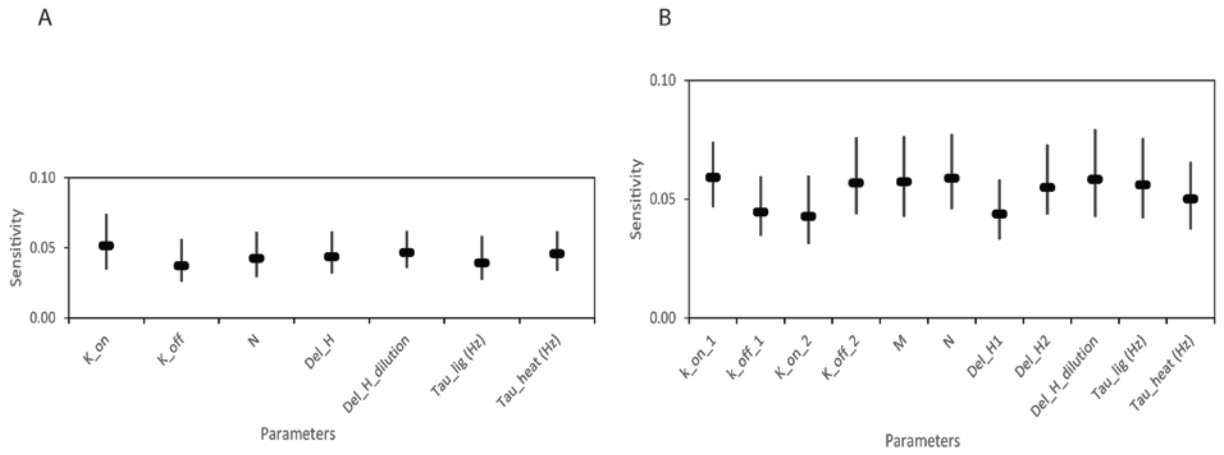


*Figure 1 The global sensitivity of fit parameters assessed through PDF based approaches for (A) 2'-CMP + RNASE and (B) BH3I-1 + hBCL$_{XL}$.*

*Table 3 Parameters used to assess the PDF based global sensitivity of fit parameters obtained for 2'-CMP + RNASE and (B) BH3I-1 + hBCL$_{XL}$ using SAFE program.*

| **Parameters** | 2'-CMP + RNASE | BH3I-1 + hBCL$_{XL}$ |
|---|---|---|
| Unconditional CDF $(NU)$ | 20 | 20 |
| Conditional CDF $(NC)$ | 20 | 20 |
| Conditional points $(n)$ | 20 | 20 |
| Kolmogorov-Smirnov statistics | 'max' (Statistics)<br>0.05 (alpha value)<br>100 (Bootstrap) | 'max' (Statistics)<br>0.05 (alpha value)<br>100 (Bootstrap) |

*Table 4 Fit parameters obtained through NDH analysis (Origin-ITC) used for simulations of thermogram via dynamic approach.*

| | RNAHH | PROTDB | PERSSON | FEOTF54 |
|---|---|---|---|---|
| **Model** | One sites | Sequential binding sites (2 sites) | Sequential binding sites (4 sites) | Two sites (independent) |
| **Parameters** | | | | |
| Equilibrium constants ($K_{eq}$) | 5.59 ± 0.13 × $10^4$ $(K_1)$ | 4.13 ± 0.20 × $10^7$ $(K_1)$<br><br>1.40 ± 0.09 × $10^5$ $(K_2)$ | 2.39 ± 0.03 × $10^3$ $(K_1)$<br><br>114 ± 1.6 $(K_2)$<br><br>2.26 ± 0.03 × $10^3$ $(K_3)$<br><br>22.3 ± 0.22 $(K_4)$ | 1.18 ± 0.40 × $10^{10}$ $(K_1)$<br><br>3.46 ± 0.91 × $10^6$ $(K_2)$ |
| Stoichiometric constant | 1.02 ± 0.00161 $(m\ index)$ | 1.0 $(m\ index; fixed)$<br><br>1.0 $(n\ index; fixed)$ | 1.0 $(m\ index; fixed)$<br><br>1.0 $(n\ index; fixed)$<br><br>1.0 $(o\ index; fixed)$<br><br>1.0 $(r\ index; fixed)$ | 1.06 ± 0.00499 $(m\ index)$<br><br>0.941 ± 0.0074 $(n\ index)$ |
| Thermodynamic constants $(\Delta H\ cals/mol)$ | -1.354 ± 0.0029 × $10^4$ $(\Delta H_{bind})$ | -8194 ± 23.9 $(\Delta H_{bind_1})$<br><br>-3121 ± 31.5 cal/mol $(\Delta H_{bind_2})$ | 314.8 ± 6.18 $(\Delta H_{bind_1})$<br><br>4769 ± 75.5 $(\Delta H_{bind_2})$<br><br>834.9± 78.6 $(\Delta H_{bind_3})$<br><br>6962± 37.8 $(\Delta H_{bind_4})$ | 767.3 ± 91.7 $(\Delta H_{bind_1})$<br><br>1.203 ± 0.134 × $10^4$ $(\Delta H_{bind_2})$ |
